# Large-Scale Comparative Transcriptomic Analysis of CHO Cell Functional Adaptation to Recombinant Monoclonal Antibody Production

**DOI:** 10.1101/2024.06.27.600995

**Authors:** Cristina N. Alexandru-Crivac, Joseph F. Cartwright, Ryan Taylor, Bernadette M. Sweeney, Marc Feary, Keerthi T. Chathoth, Daniel K. Fabian, Harry Allsopp, Adam J. Brown, David C. James

## Abstract

To comparatively evaluate cellular constraints on recombinant monoclonal antibody (mAb) production by Chinese Hamster Ovary (CHO) cells, we analysed the transcriptomes of 24 clonally derived CHO cell lines engineered with PiggyBac transposon technology to stably produce four recombinant monoclonal antibodies (mAbs) at varying specific production rates. Fed-batch cultures were sampled at exponential (day 5) and stationary (day 10) phases of culture for analysis by RNA-Seq. Recombinant mRNAs accounted for a large proportion of total mRNA across all clones, and efficient use of heavy chain (HC) mRNA to synthesise recombinant mAb (qP per HC mRNA) varied significantly with respect to both mAb product and cell line. Comparative bioinformatic analyses of CHO transcriptomes focussed on mAb specific production rate and utilised both data-driven and hypothesis-led approaches, specifically (i) production or non-production of recombinant mAb, (ii) changes in the abundance of functional groups of mRNAs abundance with mAb specific production rate and (iii) comparative analysis of informatically-mined gene subsets associated with cellular functions hypothesised to impact recombinant mAb synthesis and secretion. These analyses revealed widespread constitutive and adaptive changes in mRNA abundance associated with mAb production across a variety of cellular functions. Typically, most mechanistically consistent changes in mRNA abundance co-varying with mAb production were evident at the stationary phase sample point. These data revealed both recombinant mAb-specific limitations on cellular synthetic capacity and a generic adaptive strategy used by CHO cells to support high-level mAb production. The latter was achieved by directed and permissive regulation of endoplasmic reticulum and other processes to accommodate increased synthetic flux.

## Introduction

Chinese hamster ovary (CHO) cells are known for their efficiency in expressing correctly folded and glycosylated recombinant proteins, making them one of the most widely used cell hosts for biopharmaceutical production (Szkodny and Lee, 2022). Over the years, significant improvements in the productivity of both stable and transient production systems have been achieved, with multiple grams per litre volumetric titres now routinely achievable (Zhou et al., 2021; Schmitt et al., 2020). However, some recombinant proteins remain difficult-to-express (DTE), which is often associated with protein-specific constraints on post-translational folding, assembly and processing (Alves and Dobrowsky, 2017, Mathias et al., 2020). These bottlenecks can become increasingly problematic as the number, complexity and diversity of biologicals being produced by CHO expression systems continues to rise (Walsh and Walsh, 2022).

To increase volumetric titre of diverse recombinant proteins produced by CHO cells many genetic, vector, cell, media and process engineering strategies have been reported (Fischer et al., 2015; Tihanyi and Nyitray, 2020). A number of studies have found significant genetic and functional differences between CHO cell lineages with respect to recombinant protein production and processing (Lewis et al., 2013; Reinhart et al., 2019; Könitzer et al., 2015; Lakshmanan et al., 2019). Associated with this it is also arguable that, consequently, ‘omics analyses of CHO cell function has not yet yielded generically useful strategies for cell engineering. Multi-omics analyses of CHO cell culture systems have been used to characterise global cellular adaptations in recombinant protein producing CHO cell lines (Yusufi *et al*., 2017), and to describe the effect of cell culture conditions, such as pH on cell growth, product titre and quality (Lee *et al*., 2021). Particularly, RNAseq data analysis has been widely used to define CHO cell line-specific transcriptomic signatures (Monger *et al*., 2017, Singh *et al*., 2018), characterise cellular states of CHO cell cultures (Tossolini *et al*., 2018), investigate differential expression profiles linked with productivity and growth variants (Sha *et al*., 2018), as well as identify gene targets to reduce apoptosis (Orellana *et al*., 2021). There is, however, limited consensus between different studies (Tamošaitis and Smales, 2018) and therefore difficulties in making inferences with respect to CHO cell engineering.

Nevertheless, for all CHO cells, functional conservation does exist (and is selected for) to maintain broadly similar CHO cell phenotypes: all CHO cells exhibit Warburg metabolism (Buchsteiner *et al*., 2018), similar N-glycan processing capability, and can be adapted to grow in suspension in chemically defined media etc. It is perhaps therefore reasonable to speculate that stable recombinant protein production by any engineered CHO cell may be associated with generally conserved, coordinated changes in cell function where the engineered cell has always to reconcile two divergent biomass production objectives; the imposed burden of secreted recombinant protein production and continued cell proliferation. In this study we investigate whether (1) recombinant mAb production requires dynamic transcriptional re-programming or harnesses pre-existing transcriptional variation between cell; (2) changes in the transcriptional landscape that may enable recombinant protein production are conserved (or different) across clones and protein products, and (3) specific cellular functions are consistently enhanced to enable both CHO cell proliferation and recombinant protein production to occur simultaneously.

Here we present an extensive case study comparing the transcriptomes of twenty-four CHO cell lines derived from an industry used CHO host engineered to stably produce four different recombinant mAbs (six cell lines per mAb) each at different cell specific production rates. The four mAbs tested include both easy-to-express (ETE) and relatively difficult-to-express (DTE) mAbs, from different IgG classes (1 x IgG1 (DTE), 1 x IgG2 (DTE) and 2 x IgG4 (ETE)). We test the null hypothesis that clone- and product-specific variation in stable cell-specific production rate is primarily a function of recombinant gene transcription (i.e. cellular recombinant mRNA content) and associated recombinant mRNA dynamics (O’Callaghan et al., 2010), and that any transcriptomic variation observed is a function of clonal genetic variation unrelated to productivity. We also comparatively analysed CHO cell line performance and transcriptomes at both early (day 5) and late (day 10) stages of culture, using both data-driven and hypothesis-led analyses of differential expression to identify conserved mechanistic adaptations to cell function associated with product-specific recombinant protein production by CHO cells that inform new strategies for CHO cell and vector engineering for increased productivity.

## Materials and Methods

### CHO cell culture

CHOK1SV GS-KO^®^ host cells (GS Xceed® Gene Expression System, Lonza, UK), were cultured in suspension in CD-CHO medium (Gibco™, 10743029) containing 6mM L-Glutamine (Gibco™, 12569079) in Erlenmeyer flasks (T125ml, T250ml, T500ml) and maintained at 37°C, 140rpm at 5% CO_2_. Cells were sub-cultured every 3 – 4 days at a seeding density of 2×10^5^ viable cells mL^−1^. Cell concentration and viability were routinely measured using the Vi-CELL XR (Beckman Coulter, High Wycombe, UK).

### CHOK1SV GS-KO^®^ *transfection and cell line selection*

For each mAb (AMS002 (IgG1), AMS114 (IgG2), AMS058 (IgG4), and cB72.3 (IgG4)), a destination plasmid vector (GS PiggyBac^®^ transposon vector) encoding mAb light chain (LC), heavy chain (HC) and glutamine synthase (GS) selection marker genes was co-transfected in triplicate by electroporation (Gene Pulser Xcell™ Bio-Rad Laboratories) into exponential phase CHOK1SV GS-KO^®^ host cells with piggyBac transposase mRNA according to Lonza proprietary methods. Immediately following electroporation, triplicate electroporation cuvettes were pooled and cultured for 24 h post transfection, cells were centrifuged, and the medium replaced with a selective medium; CD-CHO supplemented with 50 μM MSX (Sigma, Poole, U.K.) without L-glutamine. Following recovery under selection pressure, cell pools were cloned using Bruker Beacon Optofluidic System to isolate clonal cell lines. Two non-producing controls that had also undergone GS^®^selection (GS^®^Nulls) were cultured alongside, as (1) a GS^®^ Null stable pool containing a randomly integrated vector that is not a PiggyBac^®^ vector (GS^®^Null_noPB, pXC17.4), and (2) three clonal cell populations containing a transposon integrated PiggyBac^®^vector (GS^®^ Null_PB, pXC18).Twenty-four clonally derived cells lines were generated, comprising of six cell lines producing one of the four mAbs, (AMS002 (IgG1), AMS114 (IgG2), AMS058 (IgG4), and cB72.3 (IgG4)).

### Measurement of mAb titre

Cell culture medium was centrifuged at 200 x g for 10 min and recombinant mAb in cell-free supernatant was measured by fluorescence polarisation using Valita^TM^ TITER rapid high-throughput IgG quantification assay microplates (Beckman Coulter,High Wycombe, U.K.), according to manufacturer’s instructions, using a SpectraMAx ID5 multimode plate reader (Molecular Devices, San Jose, U.S.A.).

### Fed-batch culture

Fed-batch cell culture was performed using Lonza proprietary methods (Lonza, UK). Cells were cultured in 30mls of Fed-batch culture medium and the Integrals of Viable Cell Concentration (IVCC) and Viable Cell Volume (IVCV) were calculated as described in Fernandez-Martell et al. (2018).Cell and biomass-specific mAb production (qP.Biomass) was calculated as described in Fernandez-Martell *et al*. (2018) and Cartwright *et al*. (2020). qP.Biomass was utilised to measure mAb production per unit cellular biomass, so as to be directly comparable to relative mRNA abundances in RNAseq datasets.

### Bioinformatic analyses of CHO cell transcriptomes

The workflows employed are summarised schematically in Figure 1.Pellets containing 5×10^6^ cells were collected at days 5 and 10 of fed-batch culture, as representative of mid-exponential (growth) and mid-stationary (production) phases, respectively. RNA extraction, mRNA library preparation (polyA enrichment) and RNA sequencing were performed by Novogene, at a sequencing depth of 50 million reads, using NovaSeq PE150, Q30 >= 85%. Both cell culture and RNA sequencing were performed in four batches (one for each mAb), and non-producing GS^®^Null cells, as well as one cB72.3-producing cell line were included in each batch, as inter-batch controls.

**Figure 1:**
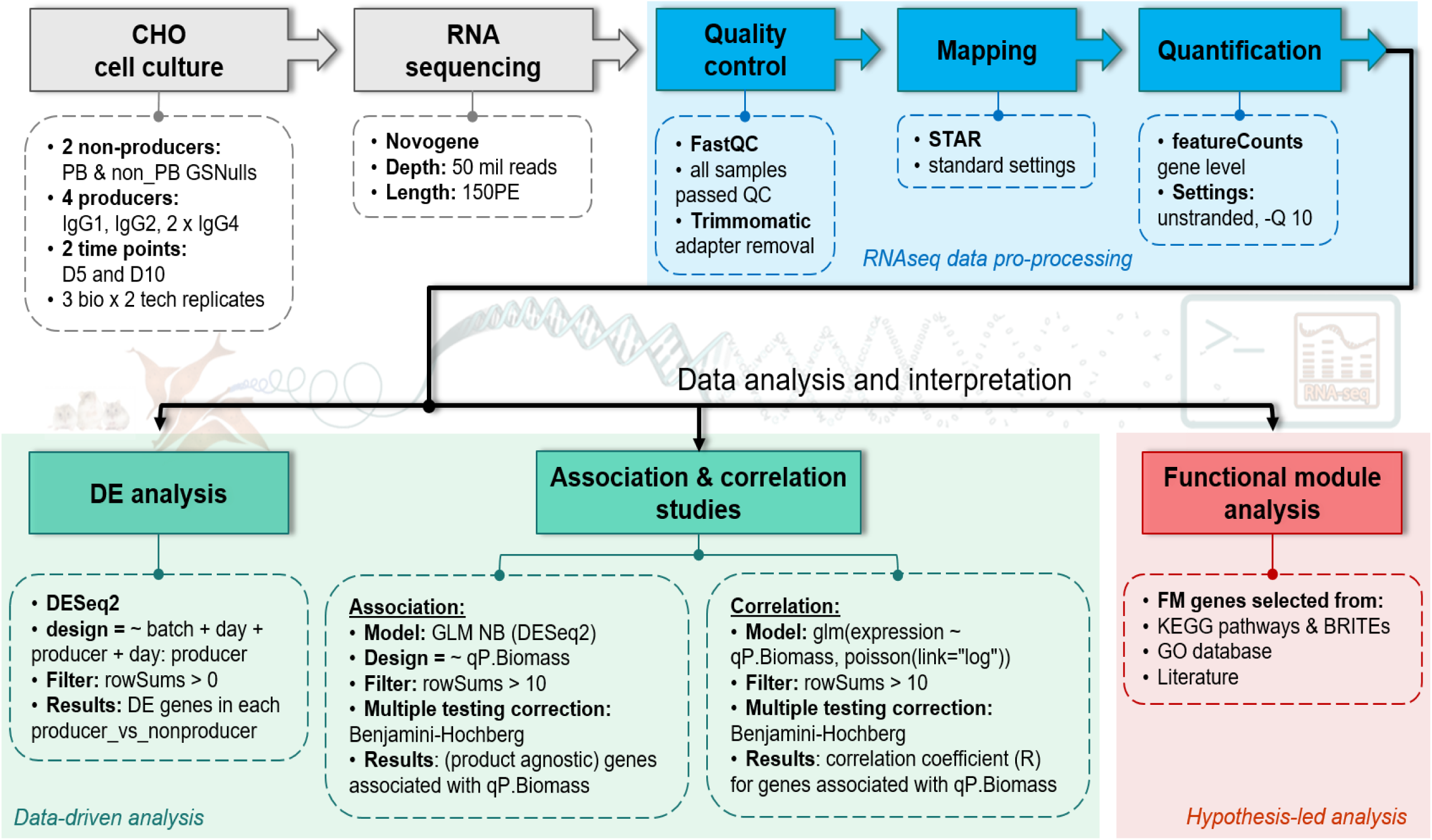
Data-driven and hypothesis-led bioinformatic analysis workflows. Cell samples from control and recombinant mAb-producing cell lines were collected at days 5 and 10 of fed-batch culture for transcriptomic analysis by RNA-Seq. Quality control, alignment of reads to the Lonza proprietary CHO genome and quantification of mRNAs was followed by (i) analyses of differential expression (DE; comparative analysis of mRNA abundance) with respect to culture sampling point and cell line specific features: mAb product, specific production rate, (ii) analyses of the degree to which mRNA abundance was associated or correlated with cell biomass specific mAb production rate and (iii) hypothesis-led comparative analyses of mRNA abundances manually assigned to curated functional modules relevant to recombinant mAb production.

RNAseq raw reads were processed through in-house modified CGAT pipelines: the quality of reads was checked with FastQC (https://www.bioinformatics.babraham.ac.uk/projects/fastqc), Novogene adapters were removed with Trimmomatic (Bolger *et al*., 2014), reads were then aligned/mapped to the Lonza proprietary reference genome (Lonza K1SV) using STAR (default settings) (Dobin *et al*., 2013), and gene-level quantification was performed with featureCounts (unstranded, -Q 10) (Liao *et al*., 2014). *DESeq2* normalised counts (Love *et al*., 2014) were used for downstream analyses and between sample comparisons, and transcripts per million (TPMs) were calculated and used in within sample comparisons. Principal component analysis (PCA) was used to identify the major sources of variation within our dataset and to visualize potential batch effects.

Two data-driven approaches were used to analyse the RNAseq dataset: differential expression (DE) analysis and association/correlation studies. DE analysis was performed with *DESeq2* package in R, between each producer *vs* non-producers, at both day 5 and day 10, and accounting for variation across batches, the two time points sampled, the different producers and the interaction between day and producer (*Expression_i_* = ∼ *batch* + *day* + *procucer* + *day*: *producer*). KEGG pathway enrichment analysis of differentially expressed genes (DEGs) was performed with *ClusterProfiler* package in R (Wu *et al*., 2021; Yu *et al*., 2012). Association and correlation studies were based on a negative binomial (NB) generalised linear model (GLM), which has been previously shown to best fit RNAseq data, as it can correctly account for the Poisson-type distribution, the overdispersion and zero inflation in counts data (Di *et al*., 2015).

Association between gene expression and bioproduction was further analysed with *DESeq2* using calculated qP.Biomass values as a continuous variable (*Expression_i_* = ∼ *qp.Biomass*). To also obtain the direction and strength of correlation between gene expression and qP.Biomass, an independent NB GLM model was run using the *glm* package in R (*glm* (*expression* ∼ *qp*.*Biomass*,*poisson*(*link* = “*log*”)). Both association and correlation analyses were performed only on mAb-producing cell lines, to correctly capture the variation and relationship between gene expression and cell-specific productivity in CHO producers. The same filtering step of lowly expressed genes (counts < 10) and Benjamin-Hochberg multiple testing correction (Haynes *et al*. 2013) were used in both *DESeq2* and the independent correlation analyses. The majority of genes were detected through both techniques, demonstrating the fitness of the model, and these commonly identified genes were used for downstream interpretation.

### Functional module analysis

Genes in fifteen functional modules (FMs) relevant to protein production were curated using annotations in the KEGG pathway, KEGG BRITE and Gene Ontology (GO) databases, as well as literature search of previous publications highlighting genes involved in key cellular processes relevant to recombinant mAb synthesis (15FMs, Table 1, Supplementary data). The number of genes in each module expressed in our dataset were recorded, as well as the percentage of genes showing significant correlation with qP.Biomass (padj <= 0.05) and the percentage of genes with strong correlation coefficients (|R| > 0.5). The differential gene expression in each producer vs non-producers, as well as the correlation with productivity were analysed and plotted for exemplar genes in functional submodules representative of the protein synthetic pathway: Protein synthesis, Competing endoplasmic reticulum (ER) secretory cargo, ER entry/translocation, ER chaperones and co-chaperones, Unfolded protein response (UPR), ER exit *via* ER-associated degradation (ERAD), ER exit from ER-to-Golgi, vesicular trafficking from Golgi-to-PM and Oxidative stress.

**Table 1:** Hypothesis-led comparative analysis of CHO cell transcriptomes: List of functional modules compared, submodules and key data source publications. The total number of genes annotated to each functional module (FM) and the number shown to be expressed in the CHO RNA-Seq datasets used in this study are indicated. Genes annotated to each FM were curated using previous literature, as well as publicly available KEGG and GO databases (Supplementary data).

| Functional module<br>(FM) | Submodules | Data Source<br>References | No. of genes |  |
| --- | --- | --- | --- | --- |
|  |  |  | Total | Expr. |
| 1. Protein synthesis | Ribosome biogenesis<br>Ribosome<br>Translation factors<br>Nucleotide metabolism | Yee *et al.* 2009,<br>Coleman *et al.* 2019,<br>Tihanyi and Nyitray, 2020,<br>Gast *et al.* 2021 | 356 | 254 |
| 2. Protein processing in the ER | Competing ER secretory cargo<br>ER entry/translocation<br>ER chaperones & co-chaperones<br>UPR | Yusufi *et al.* 2017,<br>Orellana *et al.* 2015 | 207 | 166 |
| 3. Exit from the ER | ERAD<br>ER to Golgi (incl. COPII) | Yee *et al.* 2009,<br>Orellana *et al.* 2015 | 120 | 98 |
| 4. Golgi vesicular transport | Golgi to PM<br>Cytoskeleton organisation<br>Microtubules<br>Actin | Yee *et al.* 2009,<br>Schaub *et al.* 2009,<br>Orellana *et al.* 2015 | 283 | 250 |
| 5. Glycan biosynthesis | Glycosaminoglycan biosynthesis<br>N-glycan biosynthesis<br>O-glycan biosynthesis | Hossler *et al.* 2009,<br>Park *et al.* 2017,<br>Nguyen *et al.* 2021 | 164 | 138 |
| 6. Glycan degradation | Glycosaminoglycan degradation<br>Other glycan degradation | Yusufi *et al.* 2017, | 35 | 27 |
| 7. Metabolism | AA metabolism<br>Lipid metabolism<br>FAs metabolism<br>Energy metabolism | Yee *et al.* 2009,<br>Schaub *et al.* 2009,<br>Yusufi *et al.* 2017, | 966 | 761 |
| 8. Mitochondrial metabolism | mtDNA replication<br>Import<br>Transcription & translation | Yee *et al.* 2009,<br>Schaub *et al.* 2009,<br>Zhang *et al.* 2021 | 475 | 247 |
| 9. Oxidative stress | Antioxidants<br>ROS metabolism | Yee *et al.* 2009,<br>Yusufi *et al.* 2017, | 231 | 149 |
| 10. Cell cycle | Phases<br>DNA replication<br>Checkpoints | Yee *et al.* 2009,<br>Schaub *et al.* 2009, | 163 | 151 |
| 11. Apoptosis | Pro / anti-apoptotic proteins | Chiang *et al.* 2005, Schaub<br>*et al.* 2009, | 136 | 109 |
| 12. Transcription | Transcription machinery<br>TFs | Orellana *et al.* 2015,<br>Li *et al.* 2022,<br>Kim *et al.* 2022 | 1271 | 1062 |
| 13. Transporters | ABC transporters (Import/Export)<br>AAs<br>FAs<br>Glucose<br>Mitochondrial<br>Vesicular | Geoghegan *et al.* 2018,<br>Zhang *et al.* 2021,<br>Barzadd *et al.* 2022 | 192 | 159 |
| 14. Lysosome |  | Park *et al.* 2017,<br>Barzadd *et al.* 2022 | 135 | 121 |
| 15. Spliceosome |  | Chalmers *et al.* 2017,<br>Kelsall *et al.* 2023 | 153 | 118 |

## Results

### Recombinant mAb product-specific differences in clone productivity and efficiency of recombinant mRNA utilisation

Twenty-four CHO clonal cell lines derived from the same parental cell line, comprised of four sets of six clones producing different recombinant mAbs AMS002 (IgG1), AMS114 (IgG2), AMS058 (IgG4), and cB72.3 (IgG4) were utilised for this study. Three clones derived from the same host, but lacking mAb coding sequences (GS Null) were used as controls, alongside one GS Null stable pool containing a randomly integrated vector. The clonal cell lines were selected as representatives of the range of clone productivity observed for a particular mAb.

All clones were cultured in a proprietary 12-day fed-batch process in duplicate. Clone-specific volumetric titre at the end of culture (day 12) varied between approximately 1.27 and 8.41 g L^−1^ Supplementary Table 1). Clone functional performance was characterised in terms of (i) viable cell concentration (VCD), average cell volume and the integrals of viable cell concentration and volume (IVCD and IVCV respectively, Fig. 2a) and (ii) recombinant mAb production. Volumetric mAb titre was measured at days 5 and 10 of culture when cell samples were collected for transcriptomic analysis, representative of mid-exponential and stationary phases of culture respectively (Fig. 2; Supplementary Table 1). As VCC for the selected clones varied by as much as approximately 50%, average clone specific mAb productivities were calculated as an average over days 0-10 of culture with respect to cell volume (pg (µm^3^)^−1^ day^−1^; Figs. 2c and 2d) rather than cell number (e.g. pg cell^−1^ day^−1^). Based on previous studies (e.g. Padovan-Merhar *et al*., 2015; Zhurinsky *et al*., 2010; Marguerat and Baehler, 2012; Lin and Amir, 2018) we assume a proportionate relationship between cell volume and biomass. Therefore, use of IVCV to calculate qP effectively normalises comparisons of cellular productivity across clones as per normalisation of relative mRNA abundances (e.g. *via* calculation of normalised counts).

**Figure 2:**
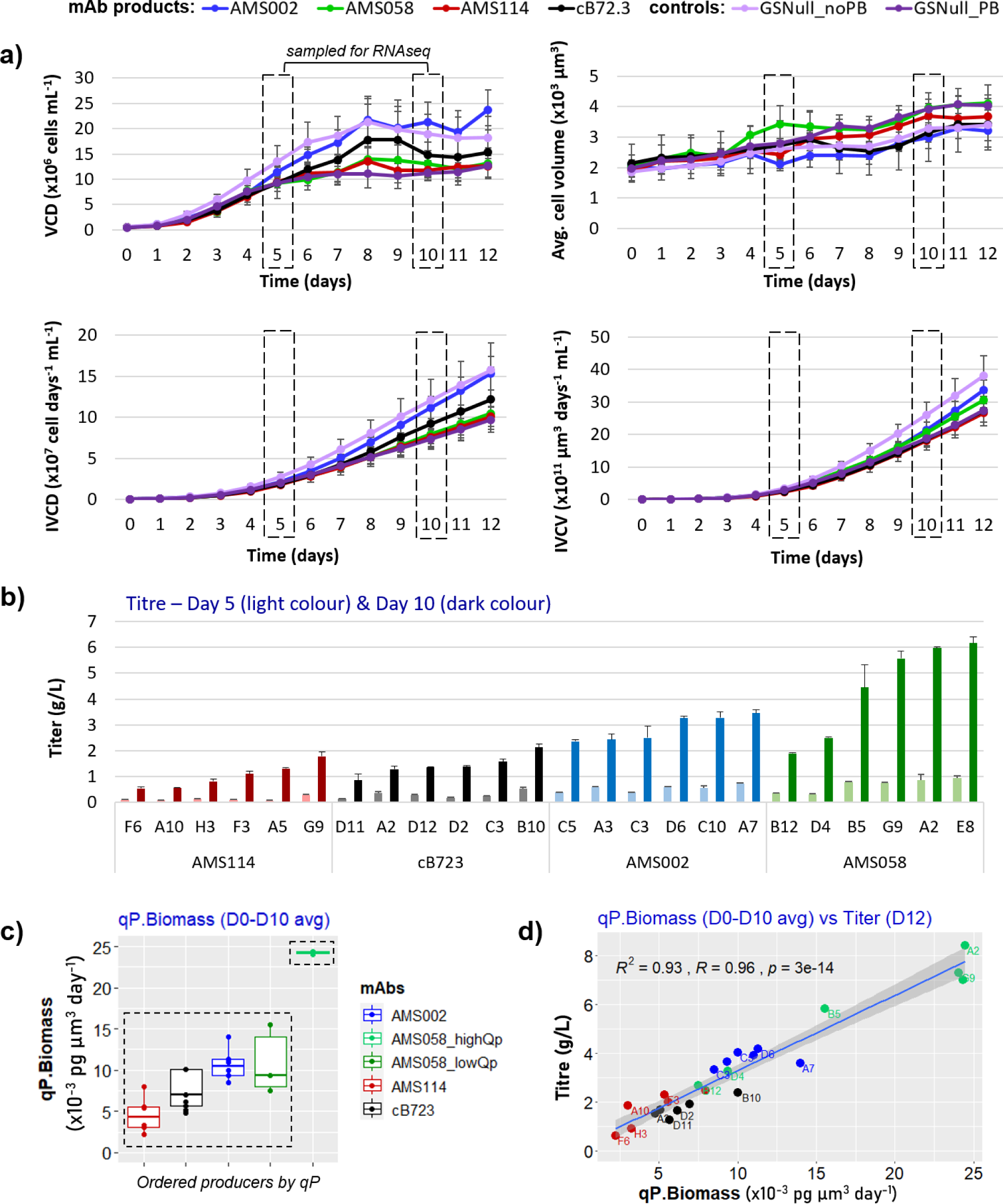
Clonally derived CHO cell lines producing four different recombinant mAbs exhibit a wide variation in productivity through fed-batch culture. Twenty-four clonally derived CHO cell lines, comprised of four sets of six clones stably producing either mAb AMS002 (IgG1), AMS114 (IgG2), AMS058 (IgG4) or cB72.3 (IgG4) were created by engineering Lonza CHOK1SVGS-KO host cells with a proprietary PiggyBac transposon vector encoding glutamine synthetase as a selection marker. Control CHO cells were engineered with both PiggyBac transposon (GSNull-PB) and randomly integrated (GSNull-noPB) vectors lacking mAb sequences. All cell lines were cultured in duplicate using a 12-day fed-batch process incorporating a transition to 33°C at day 6 of culture. **a)** Viable cell density (VCD) and average cell volume was measured daily and were used to calculate the respective integrals (IVCD and IVCV). Data for each mAb-specific group of cell lines is shown as an average ± S.D. Dashed boxes indicate the days on which cell samples were collected for transcriptomic analysis by RNA-Seq. **b)** Volumetric recombinant mAb titre for each cell line measured on days 5 and 10 of culture (n=2). **c)** Average cell biomass-specific recombinant mAb production rate across culture (day 0 to day 10) arranged as mAb product specific boxplots. Cell lines producing AMS058 were separated into low (AMS058_lowqP; cell lines B12, D4, B5) and high (AMS058_highqP; cell lines G9, A2, E8) groups. **d)** A linear regression model relating average cell line specific qP and final volumetric titre on day 12. Each point is an average of technical culture replicates (n=2) for each cell line.

As stationary phase productivity typically determines final volumetric titre (e.g. Templeton *et al*., 2013; Dickson, 2014), we confirm that across all clones qP at day 10 strongly correlated with final titre at Day 12 (Fig. 2d). Average specific productivity (days 0-10) across all clones tested varied over 10-fold between approx. 2 and pg (µm^3^)^−1^ day^−1^ (Fig. 2c) with the maximal variation in qP observed between clones producing AMS058, where three clones were designated as relatively “low producers” (AMS058_lowqP; clones B12, D4, B5) and three clones were designated as relatively “high producers” (AMS058_highqP, clones A2, E8, G9).For reference, the average qP per unit biomass observed of approx. 10 pg (µm^3^)^−1^ day^−1^ equates to approx. 33 pg cell^−1^ day^−1^. No correlation was observed between qP and any cell growth metric (VCD, IVCD, IVCV, average cell volume or biomass accumulation rate – μ; Supplementary figures S1, S2), demonstrating that there was no relationship between the capability of clones to produce recombinant mAb and accumulate cell biomass.

All mAbs used the same genetic vector components (promoters, 5’ and 3’ UTRs etc.), varying only in mAb protein sequences. Recombinant mRNA relative abundances (LC, HC and GS) were similar across all products at days 5 and 10 of culture (Fig. 3a), where GS mRNA constituted only a small proportion of total cellular mRNA (a mean of ∼0.06% at day 10) and was significantly less abundant than either HC or LC mRNA, where (for cells producing cB72.3 mAb for example) the latter reached a maximum of 39% of all mRNAs at day 10 of culture. HC and LC mRNA was proportionately less abundant relative to total mRNA at day 5 of culture. LC/HC mRNA ratio was comparable across mAbs, ranging between 2.6 −3 on both days 5 and 10 (Figure 3a).

**Figure 3:**
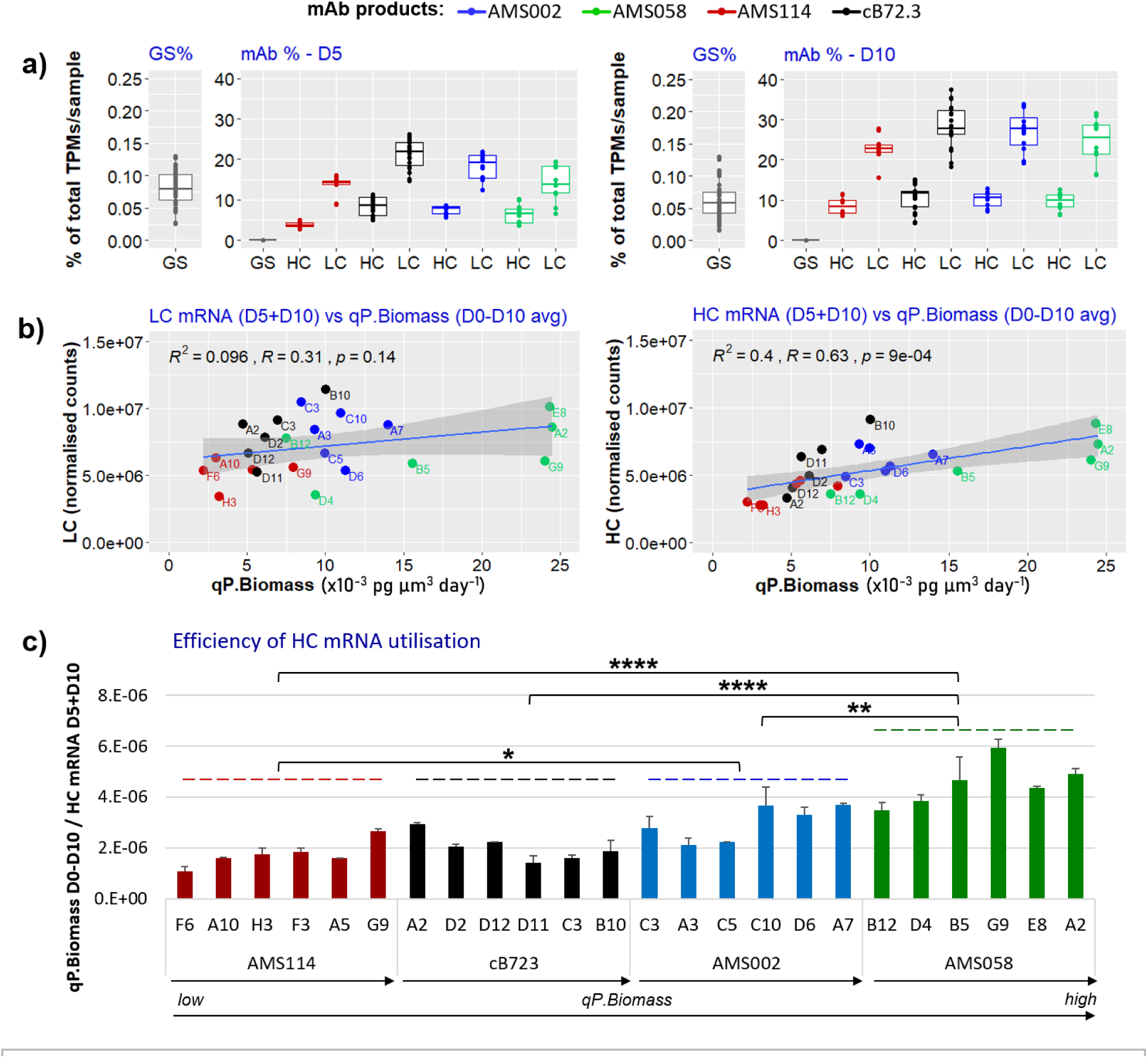
The efficiency of recombinant HC mRNA utilisation to produce mAb varies with CHO cell line and mAb product. **a)** To determine the proportion of total cellular mRNA that is recombinant mRNA the relative abundance of recombinant HC, LC and GS mRNAs were normalised to transcripts per million (TPM) for within-sample comparisons. Data are represented as mAb product-specific boxplots for HC, LC and GS mRNAs, incorporating data for each cell line (n=2). **b)** Linear regression analysis relating average culture qP (between days 0 and 10 of culture) and cumulative LC and HC mRNA abundances (ie. D5 + D10) for all cell lines (n = 2). **c)** The cell line specific efficiency of recombinant HC mRNA utilisation represented as qP per cumulative HC (D5+D10) mRNA normalised counts. Error bars represent standard deviation of the mean. Pairwise comparisons were run between averages of mRNA utilisation rates for each product (* p <= 0.05, ** p <= 0.01, *** p <= 0.001, **** p <= 0.0001).

Across all clones making the different mAbs there was a moderate but significant general correlation between recombinant mRNA abundance and qP (Figs. 3b, 3c).For three out of four recombinant mAbs there was a significant correlation between clone-specific HC mRNA abundance and average qP, although this was not apparent for clones producing AMS002. No such correlation was observed for LC mRNA. We conclude that (i) clone-specific HC mRNA abundance can significantly impact final mAb titre, and (Fig. 3b). The latter is exemplified in Figure 3c, where clone-specific variation in the efficiency of HC mRNA utilisation to synthesise mAb is represented. In general, clones producing AMS058 exhibited a HC mRNA utilisation efficiency over two-fold that of clones producing AMS114 for example. AMS058 can therefore be considered a relatively easy-to-express (ETE) mAb in comparison to more the difficult-to-express (DTE) mAb AMS114. However, even considering clones only producing AMS058 mAb for example, there was significant variation in qP/HC mRNA ratio, where 2 of 6 clones with the maximal qP (G9, A2) exhibited the highest HC mRNA utilisation efficiency. These data reveal that the extent to which cellular synthetic processes downstream of transcription constrain flux to secreted mAb can be product-specific, but there may also be a clone-specific component.

Lastly, we tested the hypothesis that mAb product- and clone-specific qP is related to the number of genes exhibiting a significant difference in mRNA abundance (differentially expressed genes, DEGs) relative to control, non-producing cells (GS Null). Our analysis revealed there is no general relationship between differences in gene expression and differences in qP across producers. (Fig. 4). Clones producing both the DTE mAb AMS114 and the ETE mAb AMS058 exhibited the lowest number of DEGs, whereas clones producing AMS002 and cB72.3 exhibited substantially more DEGs (Fig. 4, Fig. 5b). Whilst not supporting the general hypothesis that the clone-specific number of DEGs and productivity are related, these data do however reveal that the number of DEGs in CHO cells can be highly variable, from approx. 1000 DEGs (AMS114 clone F6) to approx. 10,500 DEGs (cB72.3 clone B10) (Fig. 5b). The total breadth of clonal variation in transcriptional re-programming that may be associated with recombinant mAb production is therefore large (in terms of number of genes), but not universally required to achieve maximum functional performance (AMS058).

**Figure 4:**
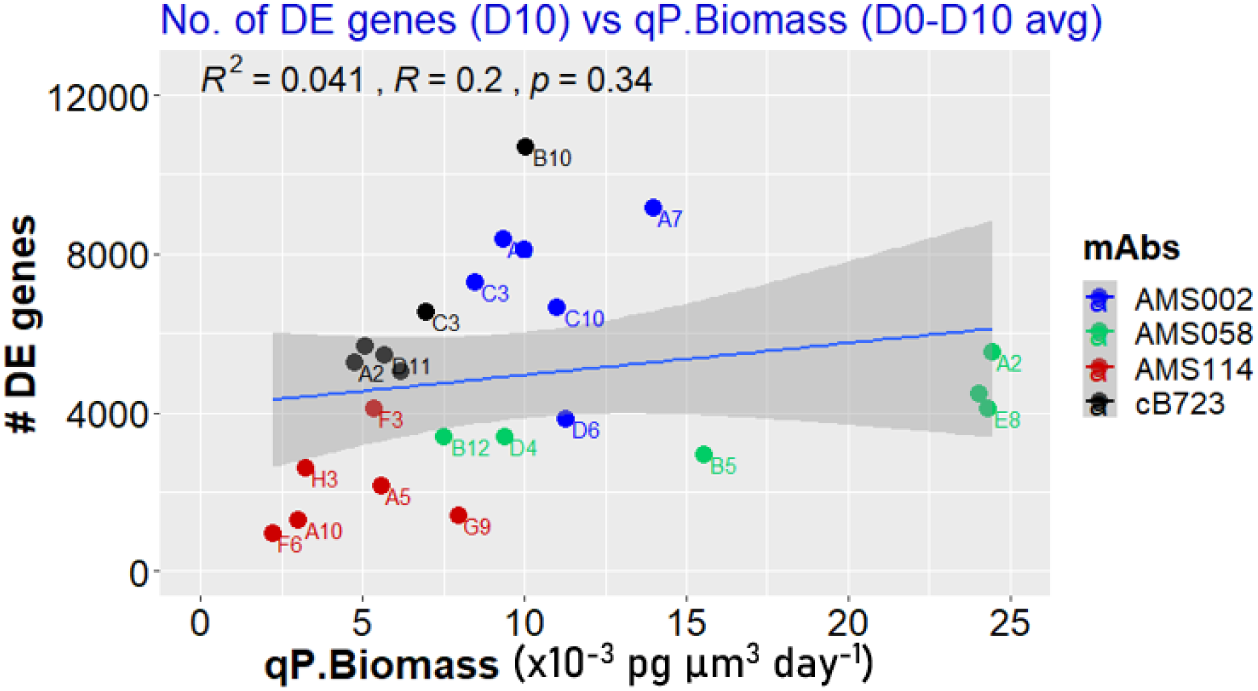
Cell line specific mAb productivity is not correlated with number of differentially expressed genes. For each cell line the number of mRNAs exhibiting a significant difference in abundance at Day 10 of culture with respect to control, non-producing (GSNull) CHO cells is plotted against the average cell biomass-specific mAb production rate through culture (n = 2). A linear regression model was fitted between average qP.Biomass and the number of differentially expressed genes observed between producers and nonproducers at day 10.

**Figure 5:**
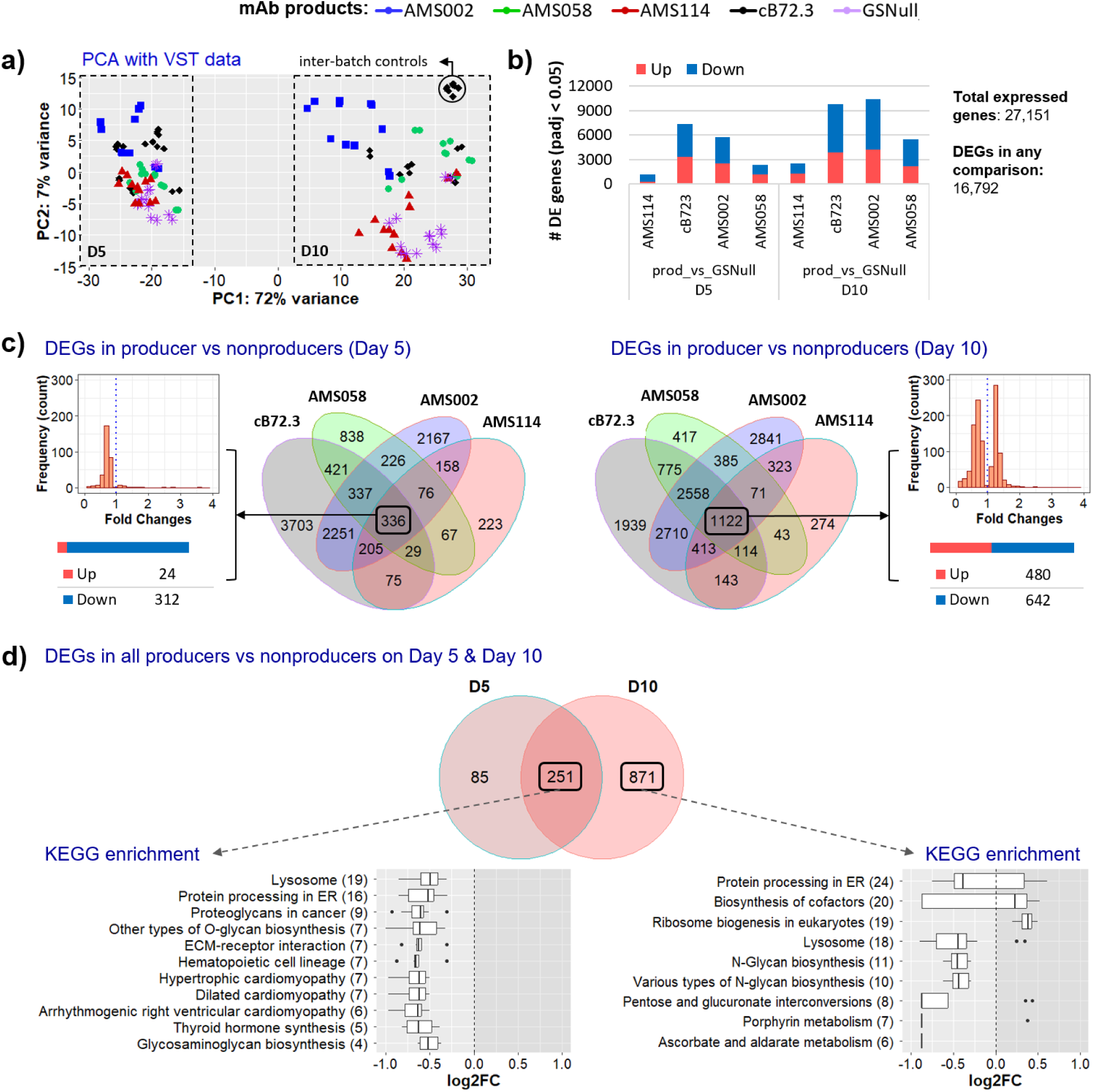
Data-driven analysis reveals functional groups of mRNAs consistently altered in abundance in mAb-producing CHO cell lines. For each cell line RNA sequencing transcript counts from days 5 and 10 of FBC were normalised using DESeq2 and the clonal distribution in gene expression was visualised using a PCA plot, excluding recombinant mRNA counts **a)**, which highlights the top two principal components: PC1 – variance by day (dashed boxes); and PC2 – variance by product (colour coded). The percentage of variance explained by each of these principal components is stated within axis labels. **b)** Number of genes that were significantly (padj ≤ 0.05) differentially expressed within all cell lines producing a given mAb product compared to GSNull controls. Bars are coloured to depict the number of upregulated (red) and downregulated (blue) genes. **c)** Four-way Venn diagrams for both days 5 and 10 of culture represent the DEGs in common between clones producing different mAbs. The number of consistent DEGs (i.e. differentially expressed within all cell lines producing all products) are in the centre of each diagram (black boxes). For each day stacked bar plots and frequency distribution plots display the number of upregulated (red) and downregulated (blue) consistent DEGs, and the range of their fold changes, respectively. **d)** The overlap of consistent DEGs on days 5 and 10 of culture. The number of constitutive DEGs (ie. differentially expressed within all clones producing all products on both days 5 and 10) are in the centre of the diagram (black box). Vertical boxplots show the variation in log2FC of genes within enriched KEGG pathways. The number of genes involved in each pathway is shown in parentheses. All overlaps were significant at P<0.05 according to Fischer’s exact test.

These data suggest that CHO cell populations are capable of utilising a broad continuum of transcriptional landscapes to support production of different mAbs. The variation in transcriptomes may possibly be generated either by selection of clonal variants with constitutive (“hardwired”) changes in gene expression (some of which may be) permissive to mAb production or adaptive (induced) changes in gene expression associated with recombinant mAb production. Taken together, our data suggests that different cellular routes to achieve recombinant mAb production are possible. For example, if we consider two clones producing different IgG4 mAbs cB72.3 (clone B10) and AMS058 (clone B12), both achieve approx. 2 g/L in fed-batch culture (Fig. 2b). However, it is clear that cB72.3 (B10) cells utilise recombinant mRNA far less efficiently to synthesise and secrete mAb than do AMS058 (B12) cells (Fig. 3c). In support of this, cB72.3 (B10) cells exhibit significantly more DEGs than AMS058 (B12) cells (∼6,800), relative to control GS^®^Null cells (Fig. 4).

### Different data-driven analyses identify functional groups of mRNAs significantly associated with efficient recombinant mAb production

To determine if specific, consistent changes in cellular mRNA abundance were generally associated with productivity (i.e. across all mAb producing clones) we initially used a data-driven approach to (i) compare the transcriptomes of non-producing (GS^®^Null) and mAb-producing cells and (ii) identify mRNAs with a change in relative abundance correlated with qP. Principal component analysis (Fig. 5a) of all transcriptomic datasets revealed that the largest (72%) source of variation in mRNA abundances across all samples was sample time point (day 5 or 10 of culture; PC1). This was followed by variation resulting from the recombinant plasmid vector being expressed (GS^®^Null, AMS002, AMS114, AMS058 or cB72.3), which was markedly smaller (PC2 – 7%). Samples were derived from four independent culture batches and the tight clustering of the inter-batch positive controls (cB72.3 clone B10) shows that there was minimal variation resulting from any batch-related effects.

### Production vs non-production: Data-driven analysis of mRNAs consistently associated with production of all mAbs

Analysis of differentially expressed genes (DEGs) revealed that, across all clones, out of a total of 27,151 CHO genes surveyed, a total of approximately 62% (16,792) exhibited a significant, consistent (i.e. in all clones producing the same mAb) change in abundance compared to GS^®^Null controls for at least one mAb at either day 5 or day 10 sample points. For clones producing different mAbs the numbers of mRNAs consistently up-or downregulated (in all clones) varied significantly. For example, on day 10, 9,781 DEGs were identified in all cB72.3 clones relative to GS^®^Null, while only 2,510 were significant in AMS114 clones (Fig. 5b). There were consistently more DEGs evident on day 10 of fed-batch culture than on day 5 for all mAb-producing clones. Across all mAbs, there was a higher proportion of downregulated (FCs between 0 and 1) than upregulated (FCs ≥ 1) mRNAs on both day 5 of FBC (56%), and day 10 (59%), relative to GS^®^Null.

Of the total number of expressed genes (27,151), 1.2% (336 genes) and 4.1% (1,122 genes) were differentially expressed by clones producing all four mAb products at days 5 and 10 respectively (Fig. 5c). These were mostly downregulated on day 5 (93%), but on day 10 approximately equal numbers (43% and 57%, respectively) of up and downregulated DEGs were observed. The FC distribution of these DEGs ranged between 0 and 7.5, with most genes showing only limited (FCs < 2) changes in expression. Of these 1,457 DEGs (day 5 or day 10), 251 were found to be differentially expressed on both days 5 and 10 in all mAb producing clones (Fig. 5d). Only 5 DEGs were upregulated relative to control across all mAb products with FCs between 1.2 and 7 (not shown, gene functions included transcription, translocation of proteins across the ER, ER-mitochondria interactions and fatty acid metabolism). The remaining 246 genes differentially expressed at both day 5 and day 10 were downregulated with FC values ranging between 0.8 and 0.008.

KEGG pathway enrichment analysis of DEGs whose collective gene expression was significantly altered in all mAb-producing CHO cells is shown in Fig. 5d, specifically with respect to (i) DEGs common to both day 5 and day 10 transcriptomes and (ii) DEGs across all producers only at day 10, where the latter may be of particular relevance to mAb production. Notably, apart from ribosome biogenesis and biosynthesis of cofactors KEGG pathways, all DEGs featured within other pathways were downregulated indicating a general loss of associated pathway function. Surprisingly, the list of downregulated pathways included protein processing in the ER – where different groups of genes being enriched either at day 5 and day 10 or only at day 10 (data not shown). There were also additional KEGG pathways that were significantly enriched in all mAb producers on only day 5 (biosynthesis of amino acids, 4 genes downregulated; lysosome, 6 genes downregulated; genes not shown).

Taken together, largely limited by the small number of DEGs associated with AMS114-producing clones, this binary analysis (i.e. production *vs* non-production) provided only a very restricted understanding of the mechanistic adaptations associated with acquisition of a mAb productive phenotype.

### Changes to the CHO transcriptional landscape correlating with cellular mAb production

To more rigorously identify genes associated with qP (Fig. 2c) we utilised an alternative means of analysis using all of our data (rather than that limited by a specific subset, eg. AMS114). A DESeq2 association analysis identified genes whose expression was significantly (padj ≤ 0.05) associated with qP, using each clone as an independent measurement of association, regardless of the recombinant mAb it was producing. This analysis does not determine the direction or strength of an association, and so a complementary correlation analysis (using a negative binomial generalised linear regression model) was applied to calculate the direction and strength of associations (correlation coefficient, R). All associated genes were also found to be significantly (padj ≤ 0.05) correlated with qP, demonstrating the fitness of the models used in both association and correlation analyses and their comparable results. Many genes were associated with qP on either day 5 only (2,379 genes), day 10 only (2,865 genes), or on both days 5 and 10 (689 genes; Fig. 6a). Most genes showed relatively weak correlations (|R| ≤ 0.5), with a maximum correlation coefficient (R) of 0.77 (Fig. 6b).

**Figure 6:**
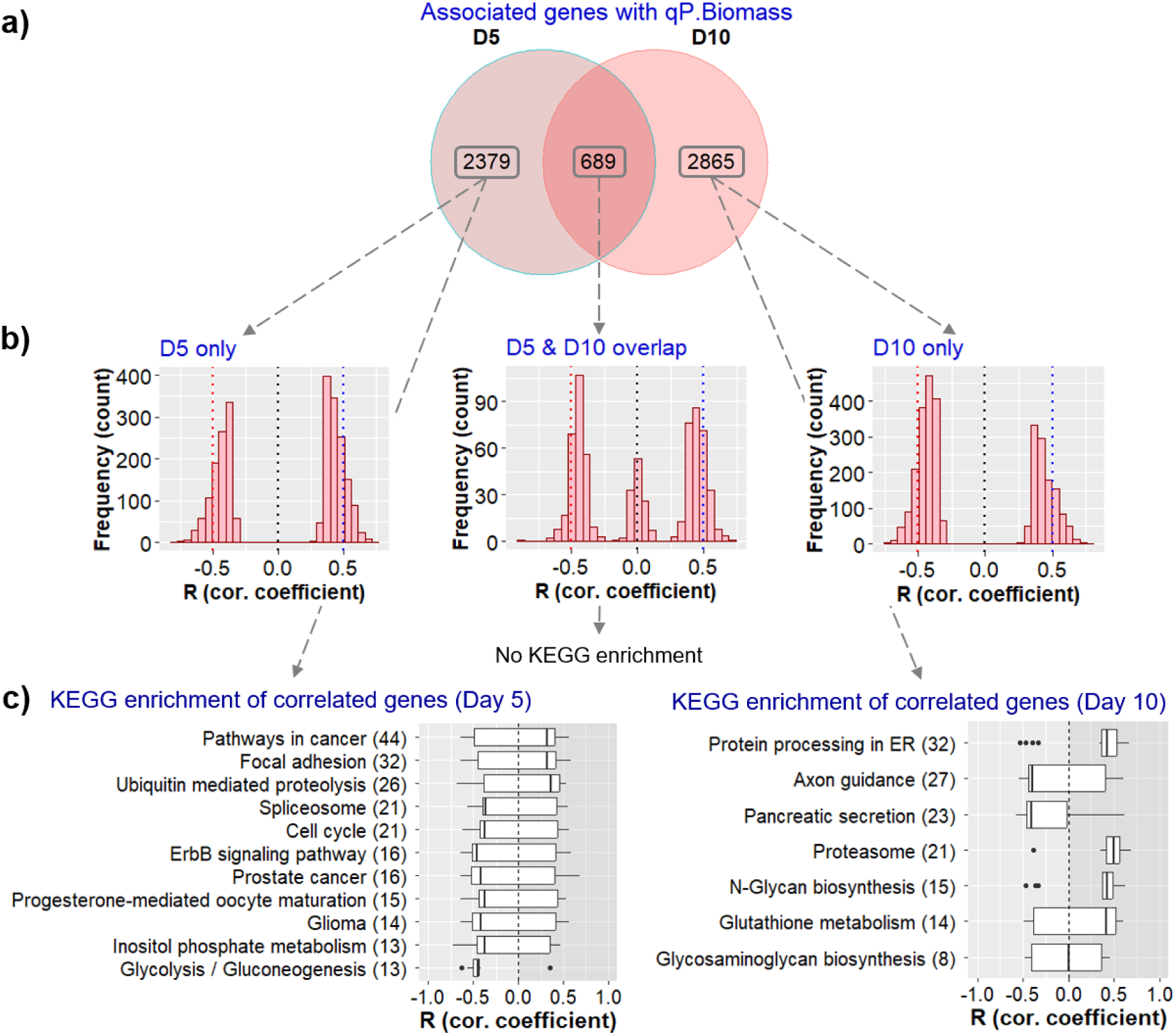
Functional groups of mRNAs co-vary with cell biomass-specific mAb production across CHO cell lines producing different recombinant mAb products. Association analysis within DESeq2 was used to identify mRNAs whose abundance is significantly (padj ≤ 0.05) associated with average qP across Days 0-5 and Days 0-10 of culture. Each cell line was used as an independent measurement of association, regardless of the recombinant mAb it was producing. A further correlation analysis (negative binomial generalised linear regression model) was applied to calculate the direction and strength of each association (correlation coefficient, R). **a)** The number of genes associated with qP on days 5 and 10 and the overlap between them. **b)** Frequency distribution plots of the correlation coefficients (R) for each associated genes display the range of strengths in correlation of gene expression with qP from each section of the Venn diagram. **c)** KEGG pathway enrichment analysis was carried out using associated genes from each section of the Venn diagram. Vertical boxplots show the variation in correlation coefficients (R) of genes within enriched KEGG pathways on days 5 and 10. The number of genes involved in each pathway is shown in parentheses. Genes associated with qP on both days 5 and 10 were not enriched in any KEGG pathways.

KEGG pathway enrichment of these genes (Fig. 6c) highlights clear differences between CHO producers on days 5 and 10. For example, on day 5 there were significant correlations within pathways related to cell growth and proliferation such as cell cycle (21 genes), glycolysis/gluconeogenesis (13 genes), and inositol phosphate metabolism (13). However, on day 10 correlations were primarily associated with pathways relevant to protein production, such as protein processing in the ER (32 genes), proteasome (21 genes), and *N*-glycan biosynthesis (15 genes). There were no enriched pathways in common between days 5 and 10. Individual genes with the strongest correlation to qP (not shown) have functions relevant to mAb production such as transcription and protein processing at different stages of the secretory pathway. Most of these genes were not previously identified through our limited mAb production vs. non-production differential expression analysis above, thus demonstrating the advantage of association/correlation studies for improving our understanding of the specific cellular mechanisms associated with mAb production.

### Hypothesis-led analyses of mAb synthetic pathway-associated mRNAs reveals a culture-phase specific hierarchy of cellular functions contributing to recombinant mAb production

Targeting specific cellular functions known to be associated with recombinant mAb production we curated gene lists for fifteen functional modules (FMs) and sub-modules *a priori* that have been reported to impact recombinant protein production. Genes were allocated to FMs using KEGG and GO database annotations, and/or previous reports within the literature of their association with recombinant protein productivity.

A top-level analysis of the 15 FMs showed the percentage of FM genes whose expression is associated with qP, both with and without a correlation R value filter (|R| ≥ 0.5), for day 5 (Fig. 7a) and day 10 (Fig. 7b). These percentages essentially indicate the extent to which each FM has been reprogrammed to adapt to mAb production. However, interpretation of relatively small FMs can be skewed by only a small number of gene associations and so the total number of genes expressed within a FM should be considered, which we have shown for reference. The percentage of FM gene associations with qP ranged between 6.9%-16.8% on day 5 and between 6.9%-34% on day 10. For gene associations where |R| ≥ 0.5 the percentage ranges were between 0%-5.4% on day 5 and between 0%-8.4% on day 10. When assessing the nature of FM gene associations with qP, we observed a higher proportion of negatively correlated genes (R < 0) on day 5 (61%, Fig. 7a), and more positively correlated genes (R > 0) on day 10 (56%, Fig. 7b). To further assess the difference between day 5 and 10 FM gene expression, we ranked FMs based upon the percentage of genes that were strongly correlated (|R| ≥ 0.5) with qP. On day 5 oxidative stress, apoptosis, metabolism, cell cycle and mitochondrial metabolism were among the top ranked FMs associated with mAb production, whereas on day 10 the top ranked FMs were protein processing in the ER, exit from the ER, lysosome, mitochondrial metabolism and Golgi vesicular transport. This accords with a biphasic transition during fed-batch culture, where cells switch from predominantly cell biomass production to recombinant protein production. Specifically referring to the day 10 functional module analysis for example (Fig. 7b) whilst these data reveal again that many FMs contribute to recombinant mAb production, they also suggest that there may be a hierarchy in the extent to which FMs impacting mAb production are re-programmed to accommodate elevated mAb production. For example, compared to FMs composed of many expressed genes (e.g. transcription, 1062 genes), few expressed genes are associated with the exit from the ER FM (98 genes). However, a much larger proportion of exit from the ER genes surveyed correlated positively with qP (19% of 98 genes, 5% with |R| ≥ 0.5) than those associated with transcription. Taken together, these data clearly demonstrate a coordinated and adaptive variation in the expression of genes associated with FMs known to control cellular processes that impact recombinant protein production.

**Figure 7:**
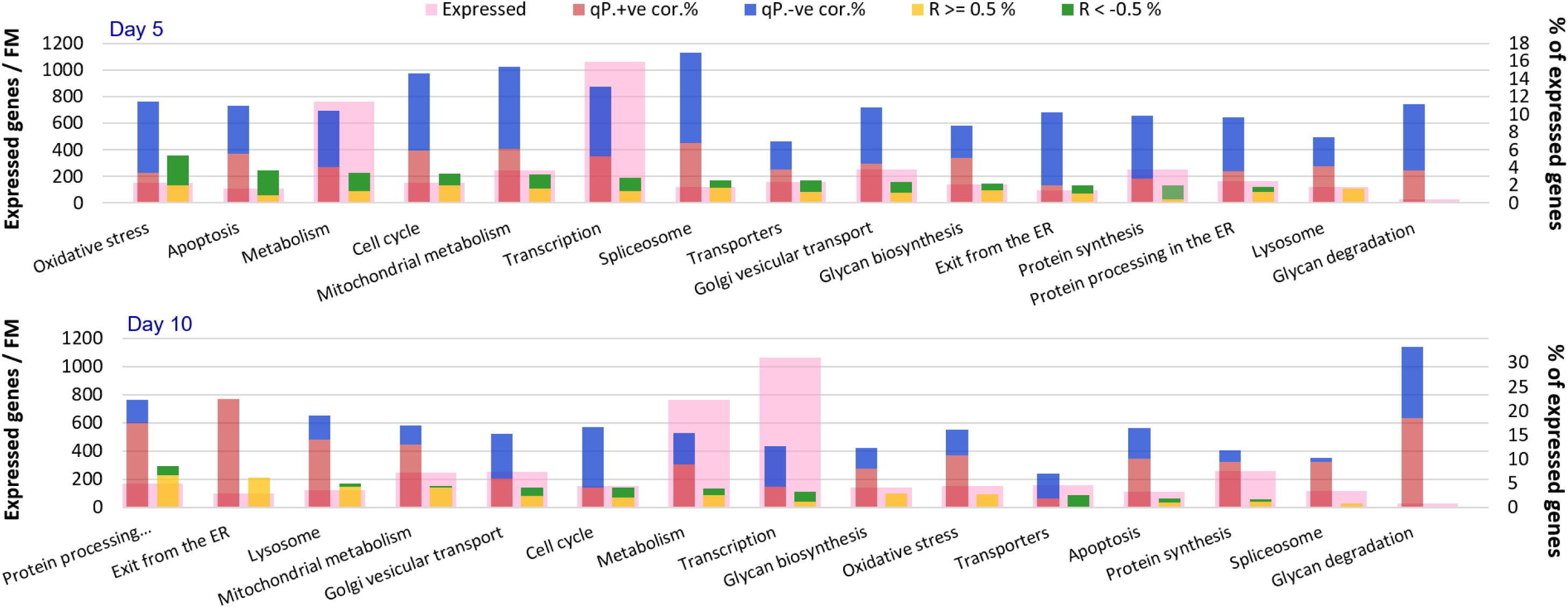
The hierarchy of cellular functional modules associated with recombinant mAb production by CHO cells varies with culture time. Key functional modules (FM) hypothesised to be associated with recombinant mAb production in CHO cells were populated by unbiased groups of genes harvested from diverse literature sources (Table 1). As described in Fig. 6, within each FM individual mRNA abundances at both the Day 5 and Day 10 culture sample points were tested for their association with biomass-specific qP (days 0-5 or days 0-10) across all CHO cell lines and mAbs. For both the Day 5 and Day 10 sample points the number of expressed genes per FM is indicated (pink bar; Table 1), with (i) the percentage of expressed genes exhibiting a positive (orange bar) or negative (blue bar) correlation with qP (padj ≤ 0.05) and (ii) the percentage of expressed genes with a positive (yellow bar) or negative (green bar) correlation coefficient equal to or greater than 0.5. For both Day 5 and Day 10 analyses, FMs are ordered from left to right with respect to the latter.

## Discussion

### General changes in recombinant mRNA abundance are associated with the maintenance of cell biomass accumulation and productivity

A fundamental difference between control and mAb-producing clones is that the latter must balance two objective functions, cell *and* mAb biomass production. Both can be considered selective pressures post-transfection although the latter is dependent on the former. In all mAb-producing clones, recombinant mAb mRNA accounted for a large proportion of total cellular mRNA (>30%). This imposition of significant recombinant mRNA synthesis on engineered cells could explain the observation that across all clones producing mAbs a large proportion of differentially expressed genes are downregulated, potentially due to competition for transcriptional activity, although these cells did not significantly differ from control, non-producing cells with respect to rate of cellular biomass accumulation. This implies that a limited downregulation of many host cell genes does not impact cellular biomass production *per se* and may represent usable capacity for recombinant gene transcription. Related to this, earlier in culture cell biomass production is predominant and reliant on maintenance of a high level of host cell protein synthesis alongside mAb synthesis. The former vastly exceeds the latter (considering total net cellular protein synthesis to include both accumulation and protein turnover parameters), even though mAb mRNA is prevalent. How is it possible to reconcile high cellular recombinant mRNA content with the maintenance of high cell biomass production? We found no evidence of an increase in protein synthetic machinery mRNAs in mAb-producing clones early in culture, although this was clearly the case later on. Previous studies have also shown that recombinant mRNA is abundant in CHO cells producing a mAb and translated as efficiently as host cell mRNAs (Kallehauge *et al*., 2017). It may be that some host cell mRNAs are preferentially recruited for translation (e.g. via the use of sequence elements such as terminal oligopyrimidine repeats characteristically associated with protein synthetic machinery mRNAs; Cockman *et al*., 2020) or other mechanisms to favour host cell mRNA translation activity such as sequestration of abundant recombinant mRNA in P-bodies (Wang *et al*., 2018). Alternatively, given that mAbs are secreted, a large proportion of mAb protein synthesised may be not measured (e.g. secretion of LC) or destined for turnover by ERAD.

We observed many productivity-related changes in host cell mRNA abundance during culture that were clearly associated with the mAb synthetic and secretory pathway (see below). However, during mid-exponential phase culture at day 5 (where cell biomass synthesis is predominant), many mRNAs associated with qP and diverse enriched KEGG pathways were downregulated and clear correlations in the abundance of mAb synthetic/secretory mRNAs with clone qP were not as widespread. In fact, the largest source (72%) of variation in mRNA abundance across our dataset was the sampling timepoint. It is well known that cells transitioning from exponential to stationary phase undergo a metabolic and physiological shift, resulting in changes to a myriad of cellular functions (Fernandez-Martell *et al*., 2017; Ahn *et al*., 2011; Rish *et al*., 2022). Indeed, this shift is evident within our analysis. For example, at day 10 of fed-batch culture, cells are in a non-proliferative state, contain more mAb mRNA, and have a higher number of genes that are differentially expressed and correlate with qP. Moreover, the genes and functional modules that are associated with qP are distinct between days 5 and 10 of fed-batch culture, shifting from generally growth-related functions to those more related to recombinant protein synthesis and secretion.

Our data therefore suggest the alteration in transcriptional programming associated with culture phase transition is a conserved response which can be modified to accommodate adaptation to recombinant mAb production during late-stage culture. The selective environment faced by engineered clones immediately post-transfection is dominated by the requirement for rapid cell proliferation minimally impacted by recombinant mAb production. This selection pressure, which favours cell biomass accumulation, persists during sub-culturing (until clone fed-batch culture performance is assessed) and clones may be expected to exhibit a degree of adaptive functional convergence, explaining the lower level of transcriptomic variability at the day 5 transcriptomic sample point. In contrast, at day 10 of culture, after the onset of stationary phase, it is likely that through genetic/functional drift (late-stage culture performance is not a selective pressure), clones should exhibit divergence in their physiological responses associated with a higher level of transcriptional variability. We have previously shown that for isolated CHO cell sub-populations late-stage culture growth performance and rapid exponential proliferation are mutually exclusive objectives (Fernandez-Martell *et al*., 2017). That many trends in mRNA abundance across different cell functions are correlated with qP are evident during late-stage culture reveals that these are a cellular response to the imposed burden of recombinant protein production rather than “hardwired” constitutive changes in gene expression. For example, it is known that cells can modulate cellular proteostasis with respect to mRNA “load”/transcriptional activity (Lin and Amir, 2018) and key determinants of protein synthetic capacity such as ribosome biogenesis and translation are co-regulated by different homeostatic mechanisms (Ni and Buszczak, 2023).

With respect to specific cellular processes, downstream of transcription a more generic constitutive adaptation associated with mAb production was evident – reduction in the relative abundance of some large mRNAs encoding homologous cell surface or secreted glycoproteins (Figure 5d, ECM-receptor interactions). Whilst expression of a range of cell surface proteins is obviously essential to support precursor transport, cellular structural integrity etc., some large, complex cell surface glycoproteins that mediate cell-cell interactions in a tissue context (e.g. laminins) may be functionally redundant with respect to CHO cell function *in vitro.* Thus, our data possibly implies that either selection of clonal variants with constitutively and selectively reduced expression of redundant secretory pathway cargo, or competition with abundant recombinant mRNA effectively “de-clutters” the secretory pathway for mAb production to some extent, effectively increasing capacity and reducing ER unfolded protein load. With respect to the latter, a relevant mechanistic linkage between recombinant protein production and cell biomass production is the mammalian unfolded protein response (UPR). It is obvious (based on our current knowledge of UPR mechanisms; Read and Schröder, 2021) to hypothesise that recombinant mAb production is associated with altered UPR signalling as selected cell lines producing mAbs do not attenuate global mRNA translation or exhibit apoptotic cell death as per the canonical acute UPR mechanism. Furthermore, we may envisage constitutive recombinant protein production as a chronic UPR stress which engineered CHO cells have to adapt to or avoid (Rutkowski and Kaufman, 2007). Perhaps counter-intuitively, a consistent feature of our data was a generally reduced constitutive abundance of mRNAs encoding ER resident polypeptides in mAb producing clones, including those which would typically be induced by ER stress via the UPR (Figure 5d, protein processing in the ER, N-glycan biosynthesis). All CHO cells engineered to produce mAbs did not exhibit an increased general abundance of mRNAs encoding the most abundant ER lumenal chaperones and foldases directly involved in mAb synthesis. However, the abundance of many mRNAs encoding potentially mAb assembly rate-limiting ER lumenal and secretory pathway components exhibited a positive correlation with productivity at day 10 of FBC (Figure 6c, Figure 7) indicating the specific importance of these components in maintenance of the most rapid rates of mAb folding and assembly -although these mRNAs still exhibited a lower abundance than control mRNA through culture in many cases expression was “recovered” in proportion to mAb productivity (Figure 6d) to offset the general constitutive reduction.

Taken together, we firstly hypothesise that “de-sensitisation” of the UPR (i.e. with an increased unfolded protein threshold) to avoid persistent, chronic UPR induction is a pre-requisite for recombinant mAb production that permits rapid cell proliferation. Mechanistically, it has been shown in other experimental models that a chronic UPR stress is associated with feedback suppression of a group of ER chaperones involved in mAb folding and assembly including Hspa5 (BiP) and Hsp90b1. Suppression was ascribed to silencing of ATF6alpha signalling and IRE1-dependent mRNA decay (Gomez and Rutkowski, 2016). Secondly, a re-programmed UPR/integrated stress response may still be active during late-stage culture that enables cells to respond to increased demand for mAb synthesis. For example, our data suggests that inducible elements of the UPR such as those induced by the multifunctional transactivator Atf4 (Neill and Masson, 2023) may still contribute to cellular capacity for increased qP without the associated induction of apoptosis.

Taken together, we suggest that even without an obvious increase in the abundance of traditional ER size/capacity and associated markers, ER <u>dynamic</u> capacity is effectively increased for recombinant mAb production via a combination of desensitised UPR, reduced endogenous cargo, and increased rates of mAb flux into and out of the ER (co-translational translocation, ERAD and anterograde transport; Figure 7).

### mAb product-specific use of available adaptive capacity in cellular synthetic processes

Most evident during late-stage culture (day 10), our analysis reveals that nearly all clones do not utilise the maximum available cellular adaptive/inducible capacity that could contribute to mAb production. Increases in the abundance of mRNAs associated with functional modules directly impacting mAb synthesis and secretion (e.g. protein synthesis, ER translocation, ER to Golgi transport, vesicular trafficking, N-glycan biosynthesis, cortical actin remodelling etc.) were most pronounced in clones with high recombinant mAb production (e.g. the relatively easy-to-express (ETE) mAb AMS058, clones E8, A2, G9).

As stated above we infer that many functionally coordinated changes in mRNA abundance correlated with increased qP (especially during late-stage culture) likely arise through cellular feedback responses to accommodate synthetic burden across the pathway from recombinant gene to secreted product rather than through constitutive changes in transcriptional landscape (i.e.. innate clonal variation in the parental population arising from inherent genome instability). Recombinant mAb-specific variation in clone qP (or more particularly the mAb-specific efficiency of recombinant HC mRNA utilisation) arises from mAb-specific restrictions in discrete, cellular synthetic processes early in the synthetic pathway (e.g. polypeptide synthesis, ER translocation, ER folding/assembly), and this likely reduces the subsequent requirement for adaptive capacity modulation to ensure that product synthetic rate is directly correlated to capacity of downstream cellular processes.

The most extreme example of this phenomenon is the general lack of adaptive changes in the transcriptional landscape of clones expressing the relatively difficult-to-express (DTE) IgG2 mAb AMS114. For clones producing this mAb, whilst the cellular content of recombinant mRNAs is still comparable to that of clones making other mAbs; the cellular efficiency of recombinant mRNA utilisation to synthesise secreted mAb is relatively low. Furthermore, other than limited constitutive changes, AMS114-producing clones do not exhibit the extensive repertoire of scaled adaptive changes in cellular mRNA abundances associated with mAb synthetic processes that are evident in clones making other mAbs, particularly during late-stage culture. Our data imply that the cellular limitation in AMS114 qP is early in the secretory pathway between transcription and ER entry/translocation. We hypothesise that for AMS114-producing clones a fundamental criterion for clonal outgrowth post-transfection required to permit maintenance of cell proliferation and proteostasis is the avoidance of ER stress via minimisation of unfolded ER lumenal recombinant polypeptides. AMS114 cell lines exhibit a reduced overall abundance of protein synthetic mRNAs specifically during late-stage culture (Figure 5d, ribosome biogenesis, Figure 7). We can find no evidence that co-translational or other ERAD processes are specifically induced in these clones. For clones producing other mAbs, which all exhibit a concerted increase in the abundance of protein synthetic mRNAs during late-stage culture, we speculate that polypeptide-specific events such as signal peptide mediated ER translocation and folding/assembly kinetics may be slower where, for example the abundance some mRNAs associated these processes scaled with product-specific qP.

### Implications for CHO cell engineering to increase productivity

Initially, recombinant gene transcriptional activity is clearly a fundamentally important parameter. Across all mAbs, HC mRNA content and qP/final titre were correlated, substantiating the findings of our previous modelling studies comparing the control exerted by discrete synthetic processes on mAb production by CHO cells through culture (O’Callaghan *et al*., 2010; McLeod *et al*., 2011). We therefore conclude that, in general, technologies to increase transcription of recombinant genes would be useful. This could be achieved directly using highly active synthetic promoters (Brown *et al*., 2017; Johari *et al*., 2019; Sergeeva *et al*., 2020) or via the use of inherently multicopy genome integration systems such as those mediated by transposons (Matasci *et al*., 2011; Balasubramanian *et al*., 2015; Ahmadi *et al*., 2017). Indirectly, whilst it has been known for many years that coding sequence optimisation is an effective strategy to increase recombinant protein and mAb expression in CHO cells (Hung *et al*., 2010) it has only recently been systematically confirmed that key optimisation parameters such as codon use bias impact mRNA stability (Forrest *et al*., 2020; Narula *et al*., 2019) as a means to increase recombinant mRNA abundance. As demonstrated by this study, it is likely that the extent to which specific recombinant proteins benefit from increased recombinant mRNA abundance is protein-specific, as for progressively more DTE proteins synthetic processes downstream of transcription will assume greater control of flux (Pybus *et al*., 2014)

Related to the above, we observed that the LC/HC mRNA ratio was relatively constant across all clones producing different mAbs. This contrasts with previous studies showing that stably transfected CHO cell populations producing recombinant mAbs deriving from random transgene integration can exhibit a variation in LC/HC mRNA ratio (O’Callaghan *et al*., 2010; Kim *et al*., 2011, Davies *et al*., 2011) and that optimal LC/HC gene expression ratio may be mAb specific (Schlatter *et al*., 2005; Pybus *et al*., 2014), which is related to mAb-specific folding and assembly dynamics. Mechanisms such as homologous recombination and epigenetic silencing are known causes of altered mAb gene expression (Yang *et al*., 2010; Kim *et al*., 2011) and transfected cell populations may essentially harness these mechanisms to enable selection and outgrowth of clonal derivatives with a more optimal LC/HC ratio (for a given mAb). It is possible that our utilisation of PiggyBac^®^ transposon technology to create stable cell lines causes mAb gene expression to be relatively impervious to these mechanisms due to transgene integration into highly transcriptionally active and stable loci. Clearly a high level of production stability is a positive outcome for manufacturing clones. However, it is possible that maintenance of the integrity of recombinant genetic constructs at transposon loci also limits the range of variant expression outcomes (see above) that may beneficially impact the rate of various mAb synthetic processes (transcription, translation, translocation etc). However, optimal product-specific vector designs could be achieved using combinations of genetic elements (e.g. synthetic promoters) that encode, for example, an optimal, product-specific stoichiometry of HC and LC gene expression (Sou *et al*., 2023).

Downstream of transcription, changes in the abundance of mRNAs significantly associated with varying qP were widespread across cellular functions but individually limited in their extent. There was a vast array of subtle changes acting in concert that were associated with CHO cell adaptation to mAb production. This reflects the general principle that alterations in cell functional phenotype may be achieved via the coordinated action of numerous small changes in transcription, epigenetics or mRNA dynamics (Ecker *et al*., 2018). Even at critical interfaces such as the ER/secretory pathway it was typical for individual mRNAs to exhibit a limited increase in abundance of 50% compared to control, even over the wide range of mAb qPs observed (>10-fold) across the diverse hierarchy of cellular functions cumulatively contribute to cell and protein-specific productivity.

Together, our data suggest that, if possible, product-specific cell engineering strategies to increase qP should also be considered alongside general cell engineering approaches, as CHO cells can evidently harness a high constitutive or induced synthetic/secretion capacity that is not influenced by variations in recombinant protein structure (e.g. transcription, vesicular transport, oxidative stress management). However, product-specific protein engineering is arguably more cumbersome to implement than cell engineering, as it requires particular knowledge of recombinant polypeptide-specific constraints on productivity (e.g. translation, translocation, folding/assembly). In this respect for example, appropriate testing of polypeptide-specific signal peptides to optimise translocation into the ER is an obvious strategy (Haryadi *et al*., 2015; O’Neill *et al*., 2023), and protein-specific engineering of the folding environment within the ER to aid expression of DTEs may require more intensive and time-consuming empirical investigation and testing to achieve a solution (Johari *et al*., 2015; Cartwright *et al*., 2020; Mathias *et al*., 2020). More generally, as high recombinant protein production involves the coordinated contribution of many cellular functional modules then host cell engineering strategies that enable genome-level (or at least multigene) reprogramming of cell functional phenotype are more attainable. This could involve modulated expression of regulators of gene expression such as transcription factors (Haredy *et al*., 2013; Berger *et al*., 2020, Torres and Dickson, 2021; Kim *et al*., 2022) and miRNAs (Raab *et al*., 2022) or overexpression of single metabolic genes that have been reported to improve the ER capacity for example (Budge *et al*., 2020). Alternatively, directed evolution of CHO cell populations using specific selective environments to create CHO host cells with superior functional characteristics may be a useful strategy (Chandrawanshi *et al*., 2020: Mistry *et al*., 2021).

In summary, we hypothesise that future cell engineering strategies will likely involve a combination of recombinant protein-specific tuning of the genetic vector used to engineer cells and host cell specific genetic/evolutionary strategies to increase the efficiency and capacity of the key cellular processes that contribute to volumetric titre (e.g. Brown *et al*., 2019). Given the intrinsic diversity of CHO genomes and client recombinant proteins, we do not anticipate that a “one-size-fits-all” solution will emerge. Rather, as is the case for many engineering systems where efficient and rapid production of a range of related products differing in features is required, the concepts and methods underpinning Product Line Engineering (Clements, 2019) may prove useful; integrating vector and cell engineering toolkits to generally increase productivity and reduce development time across a portfolio of recombinant protein products.

## Supporting information

FMs annotation and KEGG enrichment

## References

Ahmadi, S., Davami, F., Davoudi, N., Nematpour, F., Ahmadi, M., Ebadat, S., Azadmanesh, K., Barkhordari, F., & Mahboudi, F. (2017). Monoclonal antibodies expression improvement in CHO cells by PiggyBac transposition regarding vectors ratios and design. PLoS ONE, 12(6). 10.1371/journal.pone.0179902

Ahn, W. S., & Antoniewicz, M. R. (2011). Metabolic flux analysis of CHO cells at growth and non-growth phases using isotopic tracers and mass spectrometry. Metabolic Engineering, 13(5), 598– 609. 10.1016/j.ymben.2011.07.002

Alves, C.S.; Dobrowsky, T.M. (2017) Strategies and considerations for improving expression of “difficult to express” proteins in CHO cells. Methods in Molecular Biology, 1603, 1–23. DOI: 10.1007/978-1-4939-6972-2_1

Balasubramanian, S., Matasci, M., Kadlecova, Z., Baldi, L., Hacker, D. L., & Wurm, F. M. (2015). Rapid recombinant protein production from piggyBac transposon-mediated stable CHO cell pools. Journal of Biotechnology, 200, 61–69. 10.1016/j.jbiotec.2015.03.001

Barzadd, M. M., Lundqvist, M., Harris, C., Malm, M., Volk, A. L., Thalén, N., Chotteau, V., Grassi, L., Smith, A., Abadi, M. L., Lambiase, G., Gibson, S., Hatton, D., & Rockberg, J. (2022). Autophagy and intracellular product degradation genes identified by systems biology analysis reduce aggregation of bispecific antibody in CHO cells. New Biotechnology, 68, 68–76. 10.1016/j.nbt.2022.01.010

Berger, A., Le Fourn, V., Masternak, J., Regamey, A., Bodenmann, I., Girod, P. A., & Mermod, N. (2020). Overexpression of transcription factor Foxa1 and target genes remediate therapeutic protein production bottlenecks in Chinese hamster ovary cells. Biotechnology and Bioengineering, 117(4), 1101–1116. 10.1002/bit.27274

Bolger, A. M., Lohse, M., & Usadel, B. (2014). Trimmomatic: A flexible trimmer for Illumina sequence data. Bioinformatics, 30(15), 2114–2120. 10.1093/bioinformatics/btu170

Brown, A. J., Gibson, S. J., Hatton, D., Arnall, C. L., & James, D. C. (2019). Whole synthetic pathway engineering of recombinant protein production. Biotechnology and Bioengineering, 116(2), 375–387. 10.1002/bit.26855

Brown, A. J., Gibson, S. J., Hatton, D., & James, D. C. (2017). In silico design of context-responsive mammalian promoters with user-defined functionality. Nucleic Acids Research, 45(18), 10906– 10919. 10.1093/nar/gkx768

Buchsteiner, M., Quek, L.-E., Gray, P., Nielsen, L.K. (2018). Improving culture performance and antibody production in CHO cell culture processes by reducing the Warburg effect. Biotechnology and Bioengineering, 115, 2315–2327. 10.1002/bit.26724

Budge, J. D., Knight, T. J., Povey, J., Roobol, J., Brown, I. R., Singh, G., Dean, A., Turner, S., Jaques, C. M., Young, R. J., Racher, A. J., & Smales, C. M. (2020). Engineering of Chinese hamster ovary cell lipid metabolism results in an expanded ER and enhanced recombinant biotherapeutic protein production. Metabolic Engineering, 57, 203–216. 10.1016/j.ymben.2019.11.007

Cartwright, J. F., Arnall, C. L., Patel, Y. D., Barber, N. O. W., Lovelady, C. S., Rosignoli, G., Harris, C. L., Dunn, S., Field, R. P., Dean, G., Daramola, O., Gibson, S. J., Peden, A. A., Brown, A. J., Hatton, D., & James, D. C. (2020). A platform for context-specific genetic engineering of recombinant protein production by CHO cells. Journal of Biotechnology, 312, 11–22. 10.1016/j.jbiotec.2020.02.012

Chalmers, F., Van Lith, M., Sweeney, B., Cain, K., & Bulleid, N. J. (2017). Inhibition of IRE1α-mediated XBP1 mRNA cleavage by XBP1 reveals a novel regulatory process during the unfolded protein response. Wellcome Open Research, 2. 10.12688/wellcomeopenres.11764.1

Chandrawanshi, V., Kulkarni, R., Prabhu, A., & Mehra, S. (2020). Enhancing titers and productivity of rCHO clones with a combination of an optimized fed-batch process and ER-stress adaptation. Journal of Biotechnology, 311, 49–58. 10.1016/j.jbiotec.2020.02.008

Chiang, G. G., & Sisk, W. P. (2005). Bcl-xL mediates increased production of humanized monoclonal antibodies in chinese hamster ovary cells. Biotechnology and Bioengineering, 91(7), 779–792. 10.1002/bit.20551

Clements, P. C. (2019). Product Line Engineering Comes to the Industrial Mainstream. INSIGHT, 22(2), 7–14. 10.1002/inst.12241

Cockman, E., Anderson, P., & Ivanov, P. (2020). Top mrnps: Molecular mechanisms and principles of regulation. In Biomolecules (Vol. 10, pp. 1–18). MDPI AG. 10.3390/biom10070969

Coleman, O., Suda, S., Meiller, J., Henry, M., Riedl, M., Barron, N., Clynes, M., & Meleady, P. (2019). Increased growth rate and productivity following stable depletion of miR-7 in a mAb producing CHO cell line causes an increase in proteins associated with the Akt pathway and ribosome biogenesis. Journal of Proteomics, 195, 23–32. 10.1016/j.jprot.2019.01.003

Davies, S.L, McLeod, J., O’Callaghan, P.M., Pybus, L.P., Sung Y.H., Wilkinson S.J., Rance, J., Racher, A.J., Young, R.J., James, D.C. (2011) Impact of gene vector design on the control of recombinant monoclonal antibody production by CHO cells. Biotechnology Progress 27, 1689–1699. doi: 10.1002/btpr.692.

Di, Y. (2015). Single-gene negative binomial regression models for RNA-Seq data with higher-order asymptotic inference. Statistics and Its Interface, 8(4), 405–418. 10.4310/SII.2015.v8.n4.a1

Dickson, A. J. (2014). Enhancement of production of protein biopharmaceuticals by mammalian cell cultures: The metabolomics perspective. Current Opinion in Biotechnology, 30, 73–79. 10.1016/j.copbio.2014.06.004

Dobin, A., Davis, C. A., Schlesinger, F., Drenkow, J., Zaleski, C., Jha, S., Batut, P., Chaisson, M., & Gingeras, T. R. (2013). STAR: Ultrafast universal RNA-seq aligner. Bioinformatics, 29(1), 15–21. 10.1093/bioinformatics/bts635

Ecker, S., Pancaldi, V., Valencia, A., Beck, S., & Paul, D. S. (2018). Epigenetic and Transcriptional Variability Shape Phenotypic Plasticity. BioEssays, 40(2), 1700148. 10.1002/bies.201700148

Fernandez-Martell, A., Johari, Y. B., & James, D. C. (2018). Metabolic phenotyping of CHO cells varying in cellular biomass accumulation and maintenance during fed-batch culture. Biotechnology and Bioengineering, 115(3), 645–660. 10.1002/bit.26485

Fischer, S., Handrick, R., & Otte, K. (2015). The art of CHO cell engineering: A comprehensive retrospect and future perspectives. In Biotechnology Advances (Vol. 33, pp. 1878–1896). Elsevier Inc. 10.1016/j.biotechadv.2015.10.015

Forrest, M. E., Pinkard, O., Martin, S., Sweet, T. J., Hanson, G., & Coller, J. (2020). Codon and amino acid content are associated with mRNA stability in mammalian cells. PLoS ONE, 15(2). 10.1371/journal.pone.0228730

Gast, V., Campbell, K., Picazo, C., Engqvist, M., Siewers, V., & Molin, M. (2021). The yeast eif2 kinase gcn2 facilitates h2o2-mediated feedback inhibition of both protein synthesis and endoplasmic reticulum oxidative folding during recombinant protein production. Applied and Environmental Microbiology, 87(15), 1–16. 10.1128/AEM.00301-21

Geoghegan, D., Arnall, C., Hatton, D., Noble-Longster, J., Sellick, C., Senussi, T., & James, D. C. (2018). Control of amino acid transport into Chinese hamster ovary cells. Biotechnology and Bioengineering, 115(12), 2908–2929. 10.1002/bit.26794

Gomez, J. A., Thomas Rutkowski, D. (2016) Experimental reconstitution of chronic ER stress in the liver reveals feedback suppression of BiP mRNA expression. eLife, 5(DECEMBER2016). 10.7554/eLife.20390.001

Haredy, A. M., Nishizawa, A., Honda, K., Ohya, T., Ohtake, H., & Omasa, T. (2013). Improved antibody production in Chinese hamster ovary cells by ATF4 overexpression. 65, 993–1002. 10.1007/s10616-013-9631-x

Haryadi, R., Ho, S., Kok, Y. J., Pu, H. X., Zheng, L., Pereira, N. A., Li, B., Bi, X., Goh, L. T., Yang, Y., & Song, Z. (2015). Optimization of heavy chain and light chain signal peptides for high level expression of therapeutic antibodies in CHO cells. PLoS ONE, 10(2). 10.1371/journal.pone.0116878

Hossler, P., Khattak, S. F., & Li, Z. J. (2009). Optimal and consistent protein glycosylation in mammalian cell culture. In Glycobiology (Vol. 19, pp. 936–949). 10.1093/glycob/cwp079

Hung, F., Deng, L., Ravnikar, P., Condon, R., Li, B., Do, L., Saha, D., Tsao, Y. S., Merchant, A., Liu, Z., & Shi, S. (2010). mRNA stability and antibody production in CHO cells: Improvement through gene optimization. Biotechnology Journal, 5(4), 393–401. 10.1002/biot.200900192

Johari, Y. B., Brown, A. J., Alves, C. S., Zhou, Y., Wright, C. M., Estes, S. D., Kshirsagar, R., & James, D. C. (2019). CHO genome mining for synthetic promoter design. Journal of Biotechnology, 294, 1–13. 10.1016/j.jbiotec.2019.01.015

Johari, Y. B., Estes, S. D., Alves, C. S., Sinacore, M. S., & James, D. C. (2015). Integrated cell and process engineering for improved transient production of a “difficult-to-express” fusion protein by CHO cells; Integrated cell and process engineering for improved transient production of a “difficult-to-express” fusion protein by CHO cells. Biotechnology and Bioengineering, 112, 2527– 2542. 10.1002/bit.25687/abstract

Kallehauge, T. B., Li, S., Pedersen, L. E., Ha, T. K., Ley, D., Andersen, M. R., Kildegaard, H. F., Lee, G. M., & Lewis, N. E. (2017). Ribosome profiling-guided depletion of an mRNA increases cell growth rate and protein secretion. Scientific Reports, 7. 10.1038/srep40388

Kelsall, E., Harris, C., Sen, T., Hatton, D., Dunn, S., & Gibson, S. (2023). Interplay of heavy chain introns influences efficient transcript splicing and affects product quality of recombinant biotherapeutic antibodies from CHO cells. MAbs, 15(1). 10.1080/19420862.2023.2242548

Kim, M., O’Callaghan, P. M., Droms, K. A., & James, D. C. (2011). A mechanistic understanding of production instability in CHO cell lines expressing recombinant monoclonal antibodies. Biotechnology and Bioengineering, 108(10), 2434–2446. 10.1002/bit.23189

Kim, S. H., Baek, M., Park, S., Shin, S., Lee, J. S., & Lee, G. M. (2022). Improving the secretory capacity of CHO producer cells: The effect of controlled Blimp1 expression, a master transcription factor for plasma cells. Metabolic Engineering, 69, 73–86. 10.1016/j.ymben.2021.11.001

Könitzer, J.D., Müller, M.M., Leparc, G., Pauers, M., Bechmann, J., Schulz, P., Schaub, J., Enenkel, B., Hildebrandt, T., Hampel, M., Tolstrup, A.B. (2015) A global RNA-seq-driven analysis of CHO host and production cell lines reveals distinct differential expression patterns of genes contributing to recombinant antibody glycosylation. Biotechnology Journal, 10(9), 1412–1423. 10.1002/biot.201400652.

Lakshmanan, M., Kok, Y.J., Lee, A.P., Kyriakopoulos, S., Lim, H.L., Teo, G., Poh, S.L., Tang, W.Q., Hong, J., Tan, A.H., Bi, X., Ho, Y.S., Zhang, P., Ng, S.K., Lee, D.Y. (2019) Multi-omics profiling of CHO parental hosts reveals cell line-specific variations in bioprocessing traits. Biotechnology and Bioengineering, 116(9), 2117–2129. 10.1002/bit.27014

Lee, A. P., Kok, Y. J., Lakshmanan, M., Leong, D., Zheng, L., Lim, H. L., Chen, S., Mak, S. Y., Ang, K. S., Templeton, N., Salim, T., Wei, X., Gifford, E., Tan, A. H. M., Bi, X., Ng, S. K., Lee, D. Y., Ling, W. L. W., & Ho, Y. S. (2021). Multi-omics profiling of a CHO cell culture system unravels the effect of culture pH on cell growth, antibody titer, and product quality. Biotechnology and Bioengineering, 118(11), 4305–4316. 10.1002/bit.27899

Lewis, N. E., Liu, X., Li, Y., Nagarajan, H., Yerganian, G., O’Brien, E., Bordbar, A., Roth, A. M., Rosenbloom, J., Bian, C., Xie, M., Chen, W., Li, N., Baycin-Hizal, D., Latif, H., Forster, J., Betenbaugh, M. J., Famili, I., Xu, X., … Palsson, B. O. (2013). Genomic landscapes of Chinese hamster ovary cell lines as revealed by the Cricetulus griseus draft genome. Nature Biotechnology, 31(8), 759–765. 10.1038/nbt.2624

Li, Z. M., Fan, Z. L., Wang, X. Y., & Wang, T. Y. (2022). Factors Affecting the Expression of Recombinant Protein and Improvement Strategies in Chinese Hamster Ovary Cells. In Frontiers in Bioengineering and Biotechnology (Vol. 10). Frontiers Media S.A. 10.3389/fbioe.2022.880155

Liao, Y., Smyth, G. K., & Shi, W. (2014). FeatureCounts: An efficient general purpose program for assigning sequence reads to genomic features. Bioinformatics, 30(7), 923–930. 10.1093/bioinformatics/btt656

Lin, J., & Amir, A. (2018). Homeostasis of protein and mRNA concentrations in growing cells. Nature Communications, 9(1). 10.1038/s41467-018-06714-z

Manimaran, S., Selby, H. M., Okrah, K., Ruberman, C., Leek, J. T., Quackenbush, J., Haibe-Kains, B., Bravo, H. C., & Evan Johnson, W. (2016). BatchQC: Interactive software for evaluating sample and batch effects in genomic data. Bioinformatics, 32(24), 3836–3838. 10.1093/bioinformatics/btw538

Marguerat, S., & Bähler, J. (2012). Coordinating genome expression with cell size. In Trends in Genetics (Vol. 28, pp. 560–565). 10.1016/j.tig.2012.07.003

Matasci, M., Baldi, L., Hacker, D. L., & Wurm, F. M. (2011). The PiggyBac transposon enhances the frequency of CHO stable cell line generation and yields recombinant lines with superior productivity and stability. Biotechnology and Bioengineering, 108(9), 2141–2150. 10.1002/bit.23167

Mathias, S., Wippermann, A., Raab, N., Zeh, N., Handrick, R., Gorr, I., Schulz, P., Fischer, S., Gamer, M., & Otte, K. (2020). Unraveling what makes a monoclonal antibody difficult-to-express: From intracellular accumulation to incomplete folding and degradation via ERAD. Biotechnology and Bioengineering, 117(1), 5–16. 10.1002/bit.27196

Mcleod, J., O’Callaghan, P. M., Pybus, L. P., Wilkinson, S. J., Root, T., Racher, A. J., & James, D. C. (2011). An empirical modeling platform to evaluate the relative control discrete CHO cell synthetic processes exert over recombinant monoclonal antibody production process titer. Biotechnology and Bioengineering, 108(9), 2193–2204. 10.1002/bit.23146

Mistry, R. K., Kelsall, E., Sou, S. N., Barker, H., Jenns, M., Willis, K., Zurlo, F., Hatton, D., & Gibson, S. J. (2021). A novel hydrogen peroxide evolved CHO host can improve the expression of difficult to express bispecific antibodies. Biotechnology and Bioengineering, 118(6), 2326–2337. 10.1002/bit.27744

Monger, C., Motheramgari, K., Mcsharry, J., Barron, N., Clarke, C. (2017). A bioinformatics pipeline for the identification of CHO cell differential gene expression from RNA-Seq data. Methods in Molecular Biology, 1603, 169–186. doi: 10.1007/978-1-4939-6972-2_11.

Narula, A., Ellis, J., Taliaferro, J. M., & Rissland, O. S. (2019). Coding regions affect mRNA stability in human cells. 10.1261/rna

Neill, G., & Masson, G. R. (2023). A stay of execution: ATF4 regulation and potential outcomes for the integrated stress response. In Frontiers in Molecular Neuroscience (Vol. 16). Frontiers Media S.A. 10.3389/fnmol.2023.1112253

Nguyen, N. T. B., Lin, J., Tay, S. J., Mariati, Yeo, J., Nguyen-Khuong, T., & Yang, Y. (2021). Multiplexed engineering glycosyltransferase genes in CHO cells via targeted integration for producing antibodies with diverse complex-type N-glycans. Scientific Reports, 11(1). 10.1038/s41598-021-92320-x

Ni, C., Buszczak, M. (2023). The homeostatic regulation of ribosome biogenesis. Seminars in Cell and Developmental Biology, 136, 13–26. Elsevier Ltd. 10.1016/j.semcdb.2022.03.043

O’Callaghan, P. M., McLeod, J., Pybus, L. P., Lovelady, C. S., Wilkinson, S. J., Racher, A. J., Porter, A., & James, D. C. (2010). Cell line-specific control of recombinant monoclonal antibody production by CHO cells. Biotechnology and Bioengineering, 106(6), 938–951. 10.1002/bit.22769

O’Neill, P., Mistry, R. K., Brown, A. J., & James, D. C. (2023). Protein-Specific Signal Peptides for Mammalian Vector Engineering. ACS Synthetic Biology, 12(8), 2339–2352. 10.1021/acssynbio.3c00157

Orellana, C. A., Marcellin, E., Schulz, B. L., Nouwens, A. S., Gray, P. P., & Nielsen, L. K. (2015). High-antibody-producing chinese hamster ovary cells up-regulate intracellular protein transport and glutathione synthesis. Journal of Proteome Research, 14(2), 609–618. 10.1021/pr501027c

Orellana, C. A., Martínez, V. S., MacDonald, M. A., Henry, M. N., Gillard, M., Gray, P. P., Nielsen, L. K., Mahler, S., & Marcellin, E. (2021). ‘Omics driven discoveries of gene targets for apoptosis attenuation in CHO cells. Biotechnology and Bioengineering, 118, 481–490. 10.1002/bit.27548

Padovan-Merhar, O., Nair, G. P., Biaesch, A. G., Mayer, A., Scarfone, S., Foley, S. W., Wu, A. R., Churchman, L. S., Singh, A., & Raj, A. (2015). Single Mammalian Cells Compensate for Differences in Cellular Volume and DNA Copy Number through Independent Global Transcriptional Mechanisms. Molecular Cell, 58(2), 339–352. 10.1016/j.molcel.2015.03.005

Park, J. H., Jin, J. H., Lim, M. S., An, H. J., Kim, J. W., & Lee, G. M. (2017). Proteomic analysis of host cell protein dynamics in the culture supernatants of antibody-producing CHO cells. Scientific Reports, 7. 10.1038/srep44246

Pybus, L. P., Dean, G., West, N. R., Smith, A., Daramola, O., Field, R., Wilkinson, S. J., & James, D. C. (2014). Model-Directed Engineering of “‘Difficult-to-Express’” Monoclonal Antibody Production by Chinese Hamster Ovary Cells. Biotechnol. Bioeng, 111, 372–385. 10.1002/bit.25116/abstract

Raab, N., Zeh, N., Schlossbauer, P., Mathias, S., Lindner, B., Stadermann, A., Gamer, M., Fischer, S., Holzmann, K., Handrick, R., & Otte, K. (2022). A blueprint from nature: miRNome comparison of plasma cells and CHO cells to optimize therapeutic antibody production. New Biotechnology, 66, 79–88. 10.1016/j.nbt.2021.10.005

Read, A., & Schröder, M. (2021). The unfolded protein response: An overview. In Biology (Vol. 10). MDPI AG. 10.3390/biology10050384

Reinhart, D., Damjanovic, L., Kaisermayer, C., Sommeregger, W., Gili, A., Gasselhuber, B., Castan, A., Mayrhofer, P., Grünwald-Gruber, C., & Kunert, R. (2019). Bioprocessing of Recombinant CHO-K1, CHO-DG44, and CHO-S: CHO Expression Hosts Favor Either mAb Production or Biomass Synthesis. Biotechnology Journal, 14(3). 10.1002/biot.201700686

Rish, A. J., Drennen, J. K., & Anderson, C. A. (2022). Metabolic trends of Chinese hamster ovary cells in biopharmaceutical production under batch and fed-batch conditions. Biotechnology Progress, 38(1), e3220. 10.1002/btpr.3220

Rutkowski, D. T., & Kaufman, R. J. (2007). That which does not kill me makes me stronger: adapting to chronic ER stress. In Trends in Biochemical Sciences (Vol. 32, pp. 469–476). 10.1016/j.tibs.2007.09.003

Schaub, J., Clemens, C., Schorn, P., Hildebrandt, T., Rust, W., Mennerich, D., Kaufmann, H., & Schulz, T. W. (2010). CHO gene expression profiling in biopharmaceutical process analysis and design. Biotechnology and Bioengineering, 105(2), 431–438. 10.1002/bit.22549

Schlatter, S., Stansfield, S. H., Dinnis, D. M., Racher, A. J., Birch, J. R., & James, D. C. (2005). On the optimal ratio of heavy to light chain genes for efficient recombinant antibody production by CHO cells. Biotechnology Progress, 21(1), 122–133. 10.1021/bp049780w

Schmitt, M.G.; White, R.N.; Barnard, G.C. (2020) Development of a high cell density transient CHO platform yielding mAb titers greater than 2g/L in only 7 days. Biotechnology Progress, 36(6), e3047. https://publons.com/publon/10.1002/btpr.3047.

Sergeeva, D., Lee, G. M., Nielsen, L. K., & Grav, L. M. (2020). Multicopy Targeted Integration for Accelerated Development of High-Producing Chinese Hamster Ovary Cells. ACS Synthetic Biology, 9(9), 2546–2561. 10.1021/acssynbio.0c00322

Sha, S., Bhatia, H., & Yoon, S. (2018). An RNA-seq based transcriptomic investigation into the productivity and growth variants with Chinese hamster ovary cells. Journal of Biotechnology, 271, 37–46. 10.1016/j.jbiotec.2018.02.008

Singh, A., Kildegaard, H. F., & Andersen, M. R. (2018). An Online Compendium of CHO RNA-Seq Data Allows Identification of CHO Cell Line-Specific Transcriptomic Signatures. Biotechnology Journal, 13(10). 10.1002/biot.201800070

Sou, S. N., Harris, C. L., Williams, R., Kozub, D., Zurlo, F., Patel, Y. D., Kallamvalli Illam Sankaran, P., Daramola, O., Brown, A., James, D. C., Hatton, D., Dunn, S., & Gibson, S. J. (2023). CHO synthetic promoters improve expression and product quality of biotherapeutic proteins. Biotechnology Progress. 10.1002/btpr.3348

Szkodny, A.C., Lee, K.H. (2022) Biopharmaceutical manufacturing: Historical perspectives and future directions. Annual Review of Chemical and Biomolecular Engineering, 13, 141–165. 10.1146/annurev-chembioeng-092220-125832

Tamošaitis, L., & Smales, C. M. (2018). Meta-Analysis of Publicly Available Chinese Hamster Ovary (CHO) Cell Transcriptomic Datasets for Identifying Engineering Targets to Enhance Recombinant Protein Yields. Biotechnology Journal, 13(10). 10.1002/biot.201800066

Templeton, N., Dean, J., Reddy, P., & Young, J. D. (2013). Peak antibody production is associated with increased oxidative metabolism in an industrially relevant fed-batch CHO cell culture. Biotechnology and Bioengineering, 110, 2013–2024. 10.1002/bit.24858/abstract

Tihanyi, B., Nyitray, L. (2020). Recent advances in CHO cell line development for recombinant protein production. In Drug Discovery Today: Technologies (Vol. 38, pp. 25–34). Elsevier Ltd. 10.1016/j.ddtec.2021.02.003

Torres, M., & Dickson, A. J. (2021). Overexpression of transcription factor BLIMP1/prdm1 leads to growth inhibition and enhanced secretory capacity in Chinese hamster ovary cells. Metabolic Engineering, 67, 237–249. 10.1016/j.ymben.2021.07.004

Tossolini, I., López-Díaz, F. J., Kratje, R., & Prieto, C. C. (2018). Characterization of cellular states of CHO-K1 suspension cell culture through cell cycle and RNA-sequencing profiling. Journal of Biotechnology, 286, 56–67. 10.1016/j.jbiotec.2018.09.007

Walsh, G. and Walsh, E (2022). Biopharmaceutical benchmarks 2022. Nature Biotechnology, 40(12), 1722–1760. 10.1038/s41587-022-01582-x

Wang, C., Schmich, F., Srivatsa, S., Weidner, J., Beerenwinkel, N., & Spang, A. (2018). Context-dependent deposition and regulation of mRNAs in P-bodies. eLife, 7, e29815. 10.7554/eLife.29815

Wu, T., Hu, E., Xu, S., Chen, M., Guo, P., Dai, Z., Feng, T., Zhou, L., Tang, W., Zhan, L., Fu, X., Liu, S., Bo, X., & Yu, G. (2021). clusterProfiler 4.0: A universal enrichment tool for interpreting omics data. Innovation, 2(3). 10.1016/j.xinn.2021.100141

Yang, Y., Mariati, Chusainow, J., & Yap, M. G. S. (2010). DNA methylation contributes to loss in productivity of monoclonal antibody-producing CHO cell lines. Journal of Biotechnology, 147(3–4), 180–185. 10.1016/j.jbiotec.2010.04.004

Yee, J. C., Gerdtzen, Z. P., & Hu, W. S. (2009). Comparative transcriptome analysis to unveil genes affecting recombinant protein productivity in mammalian cells. Biotechnology and Bioengineering, 102(1), 246–263. 10.1002/bit.22039

Yu, G., Wang, L. G., Han, Y., & He, Q. Y. (2012). ClusterProfiler: An R package for comparing biological themes among gene clusters. OMICS A Journal of Integrative Biology, 16(5), 284–287. 10.1089/omi.2011.0118

Yusufi, F. N. K., Lakshmanan, M., Ho, Y. S., Loo, B. L. W., Ariyaratne, P., Yang, Y., Ng, S. K., Tan, T. R. M., Yeo, H. C., Lim, H. L., Ng, S. W., Hiu, A. P., Chow, C. P., Wan, C., Chen, S., Teo, G., Song, G., Chin, J. X., Ruan, X., … Lee, D. Y. (2017). Mammalian Systems Biotechnology Reveals Global Cellular Adaptations in a Recombinant CHO Cell Line. Cell Systems, 4(5), 530–542.e6. 10.1016/j.cels.2017.04.009

Zhang, H. Y., Fan, Z. L., & Wang, T. Y. (2021). Advances of Glycometabolism Engineering in Chinese Hamster Ovary Cells. In Frontiers in Bioengineering and Biotechnology (Vol. 9). Frontiers Media S.A. 10.3389/fbioe.2021.774175

Zhou, H., Fang, M., Zheng, X., Zhou, W. (2021) Improving an intensified and integrated continuous bioprocess platform for biologics manufacturing. Biotechnology and Bioengineering, 118, 3618–3623. 10.1002/bit.27768

Zhurinsky, J., Leonhard, K., Watt, S., Marguerat, S., Bähler, J., & Nurse, P. (2010). A coordinated global control over cellular transcription. Current Biology, 20(22), 2010–2015. 10.1016/j.cub.2010.10.002

