## Supplementary tables and figures for "Large-Scale Comparative Transcriptomic Analysis of CHO Cell Functional Adaptation to Recombinant Monoclonal Antibody Production"

**– Supplementary data –**

Cristina N. Alexandru-Crivac ^1^, Joseph F. Cartwright^1^, Ryan Taylor^1^, B M Sweeney^3^, Marc Feary^2^, Keerthi T Chathoth^2^, Daniel K Fabian^2^, Adam J. Brown^1^, David C. James^1^*

^1^ Department of Chemical and Biological Engineering, University of Sheffield, Mappin St., Sheffield, S1 3JD, U.K.

^2^ Lonza Biologics plc, Cambridge CB10 1XL, U.K.

^3^ Lonza Biologics Plc, Slough SL1 4DX, U.K.

*To whom correspondence should be addressed.

**Keywords**

Recombinant protein, monoclonal antibody, CHO cells, bioproduction, transcriptomics.

|  | Clone | Type | Avg. cell diameter *(μm)* | Max VCD *(x10^6^ cells/mL)* | IVCD (D12)  *(x10^7^ cell days^-1^ mL^-1^)* | IVCV (D12) *(x10^12^μm^3^ days^-1^ mL^-1^)* | Titre (D12) *(g/L)* | qP.Biomass  (D0-D12) *(x10^-3^ pg μm^3^ day^-1^)* |
| --- | --- | --- | --- | --- | --- | --- | --- | --- |
| GSNull pool | 1 | GSNull_noPB | 17.02 | 21.37 | 15.75 | 4.40 | N/A | N/A |
| GSNull clones | 1 | GSNull_PB | 17.73 | 11.80 | 9.20 | 2.94 | N/A | N/A |
|  | 2 |  | 18.22 | 11.95 | 8.39 | 2.99 | N/A | N/A |
|  | 3 |  | 17.73 | 15.55 | 11.51 | 3.76 | N/A | N/A |
| AMS002 | A3 | IgG1 | 16.37 | 24.85 | 16.15 | 4.05 | 3.65 | 8.93 |
|  | A7 |  | 16.81 | 21.90 | 13.85 | 3.71 | 3.59 | 9.85 |
|  | C10 |  | 16.52 | 25.40 | 16.87 | 4.24 | 3.91 | 9.21 |
|  | C3 |  | 16.48 | 30.80 | 18.22 | 4.56 | 3.34 | 7.54 |
|  | C5 |  | 16.97 | 19.50 | 12.40 | 3.49 | 4.03 | 11.54 |
|  | D6 |  | 17.22 | 20.85 | 14.58 | 4.26 | 4.16 | 9.78 |
| AMS114 | A10 | IgG2 | 17.90 | 10.55 | 8.01 | 2.64 | 1.86 | 7.05 |
|  | A5 |  | 17.19 | 15.40 | 11.08 | 3.28 | 2.00 | 6.13 |
|  | F3 |  | 17.43 | 11.90 | 9.31 | 2.79 | 2.31 | 8.28 |
|  | F6 |  | 17.52 | 15.95 | 10.97 | 3.44 | 0.62 | 1.81 |
|  | G9 |  | 17.31 | 13.90 | 10.23 | 3.05 | 2.46 | 7.85 |
|  | H3 |  | 17.53 | 15.90 | 10.89 | 3.55 | 0.92 | 2.54 |
| AMS058 | A2 | IgG4 | 17.57 | 15.25 | 10.69 | 3.38 | 8.41 | 24.92 |
|  | B12 |  | 19.02 | 12.20 | 9.44 | 3.56 | 2.69 | 7.62 |
|  | B5 |  | 17.17 | 19.55 | 13.71 | 4.03 | 5.83 | 14.45 |
|  | D4 |  | 17.66 | 15.40 | 11.43 | 3.82 | 3.27 | 8.58 |
|  | E8 |  | 18.16 | 12.60 | 9.80 | 3.37 | 6.99 | 20.81 |
|  | G9 |  | 18.93 | 11.15 | 7.70 | 3.22 | 7.29 | 22.44 |
| cB72.3 | A3 | IgG4 | 18.15 | 17.55 | 11.60 | 3.68 | 1.53 | 4.16 |
|  | B10 |  | 18.62 | 12.68 | 8.98 | 3.19 | 2.39 | 8.29 |
|  | C3 |  | 15.28 | 25.05 | 16.38 | 3.18 | 1.92 | 6.02 |
|  | D11 |  | 15.68 | 15.00 | 10.15 | 2.22 | 1.27 | 5.80 |
|  | D12 |  | 15.41 | 29.55 | 18.58 | 3.78 | 1.68 | 4.46 |
|  | D2 |  | 15.07 | 27.00 | 16.75 | 3.18 | 1.64 | 5.15 |

**Supplementary table 1. List of CHO non-producing controls and mAb-producing clones, the type of mAb, average cell diameter, max VCD achieved, IVCD, IVCV, Titer and qP.Biomass measurements on D12.**

**Supplementary figure S1: Relationship between qP.Biomass and VCD, average cell volume or biomass accumulation rate.**

Twenty-four CHO clones, comprising of four sets of six clones producing each of the mAbs AMS002 (IgG1), AMS114 (IgG2), AMS058 (IgG4), and cB72.3 (IgG4) were cultured in duplicate within a 12-day fed-batch process alongside non-producing control cells that had also undergone GS selection (GSNulls). A linear regression model was fitted to represent the relationship between qP.Biomass and **a)** VCD values measured at D12; **b)** average cell volume measured at D12; **c)** biomass accumulation rate (u.Biomass) measured between D0 and D5, or D0 and D10. Points on the scatter plots represent the average between technical replicates (n=2) of each of the six clones producing each product.

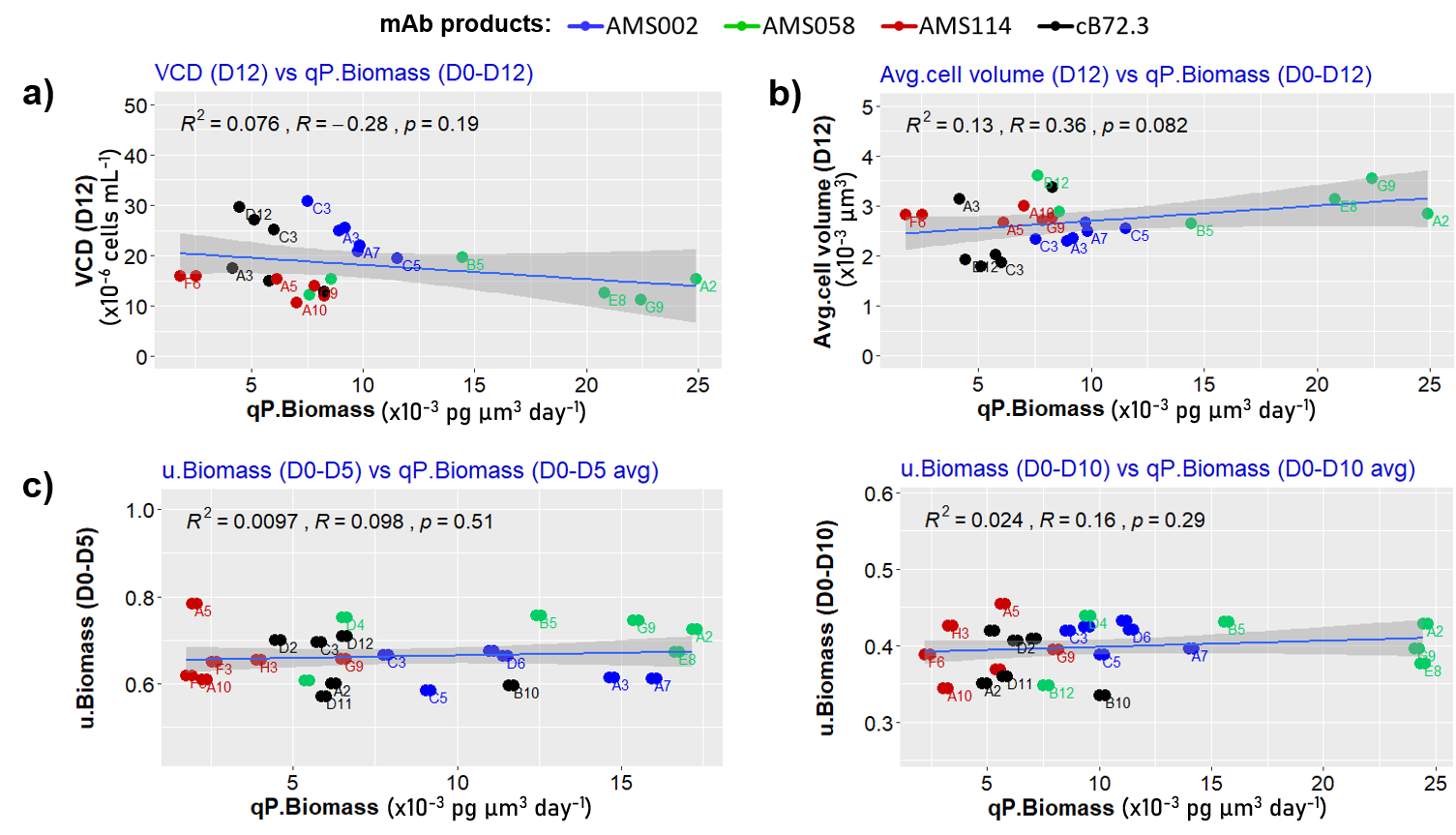

**Supplementary figure S2: Relationship between qP.Biomass and IVCD or IVCV.**

Twenty-four CHO clones, comprising of four sets of six clones producing each of the mAbs AMS002 (IgG1), AMS114 (IgG2), AMS058 (IgG4), and cB72.3 (IgG4) were cultured in duplicate within a 12-day fed-batch process alongside non-producing control cells that had also undergone GS selection (GSNulls). A linear regression model was fitted to represent the relationship between qP.Biomass and **a)** IVCD or **b)** IVCV values measured at D5, D10 and D12, respectively. Points on the scatter plots represent the average between technical replicates (n=2) of each of the six clones producing each product.

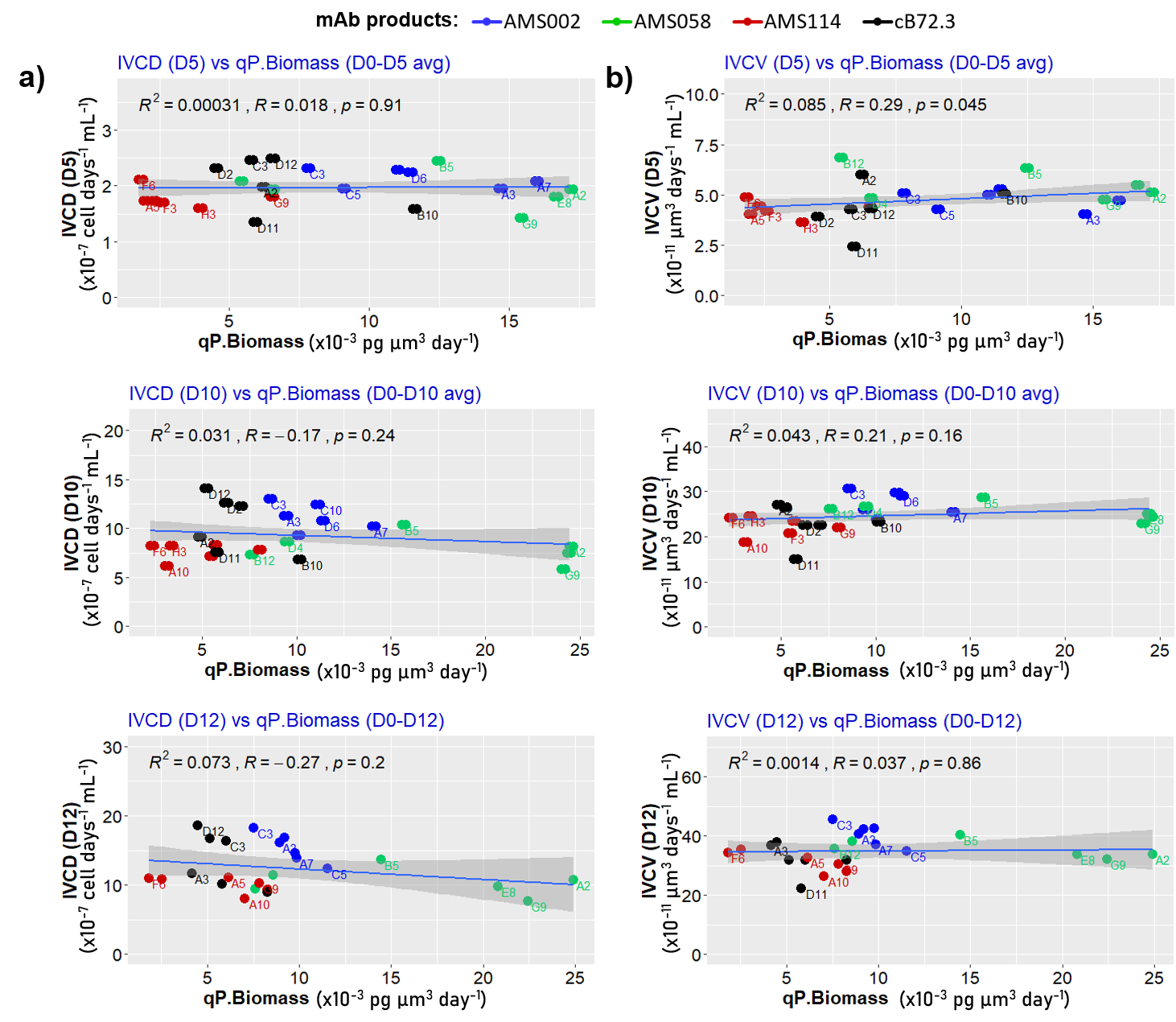
